# Ribonuclease activity undermines immune sensing of naked extracellular RNA

**DOI:** 10.1101/2024.04.23.590771

**Authors:** Mauricio Castellano, Valentina Blanco, Marco Li Calzi, Bruno Costa, Kenneth Witwer, Marcelo Hill, Alfonso Cayota, Mercedes Segovia, Juan Pablo Tosar

## Abstract

The plasma membrane and the membrane of endosomal vesicles are considered physical barriers preventing extracellular RNA uptake. While naked RNA can be spontaneously internalized by certain cells types, functional delivery of naked RNA into the cytosol has been rarely observed. Here we show that extracellular ribonucleases, mainly derived from cell culture supplements, have so far hindered the study of extracellular RNA functionality. In the presence of active ribonuclease inhibitors (RI), naked bacterial RNA is pro-inflammatory when spiked in the media of dendritic cells and macrophages. In murine cells, this response mainly depends on the action of endosomal Toll-like receptors. However, we also show that naked RNA can perform endosomal escape and engage with cytosolic RNA sensors and ribosomes. For example, naked mRNAs encoding reporter proteins can be spontaneously internalized and translated by a variety of cell types, in an RI-dependent manner. In vivo, RI co-injection enhances the activation induced by naked extracellular RNA on splenic lymphocytes and myeloid-derived leukocytes. Furthermore, naked extracellular RNA is inherently pro-inflammatory in ribonuclease-poor compartments such as the peritoneal cavity. Overall, these results demonstrate that naked RNA is bioactive and does not need encapsulation inside synthetic or biological lipid vesicles for functional uptake, making a case for nonvesicular extracellular RNA-mediated intercellular communication.

## INTRODUCTION

Lipid bilayers are considered to be impermeable to negatively charged RNA molecules. However, cells can pack and release RNA inside biological lipid nanoparticles called extracellular vesicles (EVs) (Valadi et al. 2007; Ratajczak et al. 2006; Skog et al. 2008). These vesicles can be internalized by recipient cells enabling the intercellular exchange of genetic information (Kalluri and LeBleu 2020; O’Brien et al. 2020). Similarly, therapeutic delivery of messenger RNA (mRNA) or double-stranded small interfering RNAs (siRNA) also requires RNA encapsulation inside synthetic lipid nanoparticles (LNPs).

The spontaneous uptake of naked RNA molecules is less studied and considered highly inefficient. Nevertheless, some pharmaceutical preparations containing antisense oligonucleotides (ASOs) lack cationic lipids or any other transfection reagents (Levin 2019). These drugs manage to enter cells and gain access to the cytosol or even the nucleus through an endocytosis-dependent naked RNA-specific uptake process referred to as “gymnosis” (Stein et al. 2010). Although the molecular mechanism(s) responsible for this functional uptake route are still obscure, gymnosis might depend on the increased hydrophobicity conferred by phosphorothioate bonds, a frequent modification in therapeutic ASOs (Dowdy 2017).

Gymnotic uptake and subsequent translation of naked mRNA vaccines has been reported, usually following intramuscular, intradermal or intralymphatic administration in mice (Wolff et al. 1990; Kreiter et al. 2010; Diken et al. 2011; Selmi et al. 2016). These studies have also shown that dendritic cells are the main cell type responsible for gymnotic mRNA uptake, probably through a macropinocytosis-dependent mechanism. However, intranodal injection of naked mRNA vaccines in humans failed to show a response compared with placebo (Bitounis et al. 2024). The clinical success of LNPs as delivery vectors for mRNAs and siRNAs (Akinc et al. 2019; Hou et al. 2021) has consolidated the view that cationic or ionizable lipids are required for efficient RNA uptake, as supported by analysis of delivery methods in clinical trials (Bitounis et al. 2024).

Compared with lipofection, gymnotic uptake is considered a slower and less efficient process (Stein et al. 2010). Currently, its study is mostly restricted to the context of short and highly modified therapeutic ASOs. Several factors explain the disadvantage of this uptake route for longer or unmodified RNAs. For example, endosomal escape, the rate-limiting step in RNA therapeutics (Dowdy 2023), is facilitated by an LNP-induced endosomal rupture mechanism that would not apply in the case of gymnosis. Immunological studies also argue against extracellular naked RNAs being functional. For instance, transfection of bacterial RNAs using cationic lipids triggers immune cell activation in human and murine macrophages (Sha et al. 2014; Vanaja et al. 2014), but this response is abrogated when the RNA is spiked into the media in the absence of lipofection reagents (Sha et al. 2014).

Finally, naked RNA is highly unstable in biofluids and extracellular samples, which tend to contain high quantities of active ribonucleases (RNases). This fact, coupled to the low efficiency of gymnotic uptake (Stein et al. 2010), and the apparent lack of immune cell activation induced by naked extracellular bacterial RNAs (Sha et al. 2014), explains why RNA-mediated intercellular communication is mostly—and almost exclusively—studied in the context of EVs (Tosar et al. 2021).

We have recently shown that some RNAs are intrinsically stable in the extracellular space and act as stable reservoirs of shorter RNA fragments that can enter cells spontaneously (Costa et al. 2023). In addition, when adding a broad-range ribonuclease inhibitor (RI) to human cancer cell-conditioned medium, we stabilized a previously unknown population of extracellular ribosomes that induced dendritic cell maturation (Tosar et al. 2020). These results suggest that extracellular, nonvesicular RNAs, comprising both naked RNAs and ribonucleoprotein complexes, can interact with recipient cells and induce downstream signaling responses. Importantly, to observe any functional interaction involving nonvesicular exRNAs, strict control of extracellular RNase activity is required. This is impossible in typical cell culture experiments, in which 10% fetal bovine serum (FBS) is usually employed. We hypothesize that failure to control for serum-derived RNase activity has so far precluded the study of intercellular communication pathways mediated by nonvesicular RNAs, hence underestimating the efficiency of gymnosis as a functional and efficient RNA uptake process.

Here, using a broad-spectrum ribonuclease inhibitor (RI), we show that both single-stranded and double-stranded long and short RNAs can be spontaneously internalized by different murine and human cell types in the absence of any transfection reagents. Internalized RNAs can be sensed by Toll-like receptors (TLRs) inside endosomal compartments. Furthermore, at least some RNA molecules can escape the endosomes into the cytosol and regulate or modify gene expression in recipient cells. Intravenous administration of RI enhances the inflammatory responses triggered by naked exRNA in vivo. In contrast, compartments with low RNase activity such as the peritoneal cavity seem to be inherently prone to nonvesicular exRNA recognition. Altogether, these results suggest that naked or nonvesicular exRNAs and extracellular RNases might work as antagonistic inducers and regulators of compartment-specific inflammatory responses.

## RESULTS

### Naked bacterial exRNA activates BMDCs in an RI-dependent manner

Earlier studies showed lack of exRNA recognition by innate immune cells when bacterial RNA, a well-known proinflammatory molecule, was spiked into the medium instead of transfected (Sha et al. 2014; Oldenburg et al. 2012). This suggests that the spontaneous uptake of naked exRNA is inefficient; possibly due to the fact that cells are refractory to internalizing negatively-charged nucleic acids (Dowdy 2017). Alternatively, it could simply be a consequence of RNA stability. Because cell culture experiments are typically done in the presence of FBS, a rich source of RNases, we hypothesized that this could explain the apparent lack of naked exRNA bioactivity (Tosar et al. 2020, 2021; Costa et al. 2023). To study this, we worked with primary cultures of murine bone marrow-derived cells, differentiated into antigen presenting cells (Helft et al. 2015; Na et al. 2016). We incubated these BMDCs with purified *Escherichia coli* RNA in the absence of any transfection reagents, but adding either a broad-range extracellular RNase inhibitor (+RI) or a thermally inactivated inhibitor (+RIΔ), lacking any protective activity (**Figure S1A**). Consistent with our hypothesis, naked bacterial exRNA induced a highly proinflammatory transcriptional signature only in the presence of RI (**Figure 1A-1C** and **S2**). Of note, BMDC-derived self-RNA was immunologically silent regardless of RI addition, in agreement with previous studies (Karikó et al. 2005).

**Figure 1.**
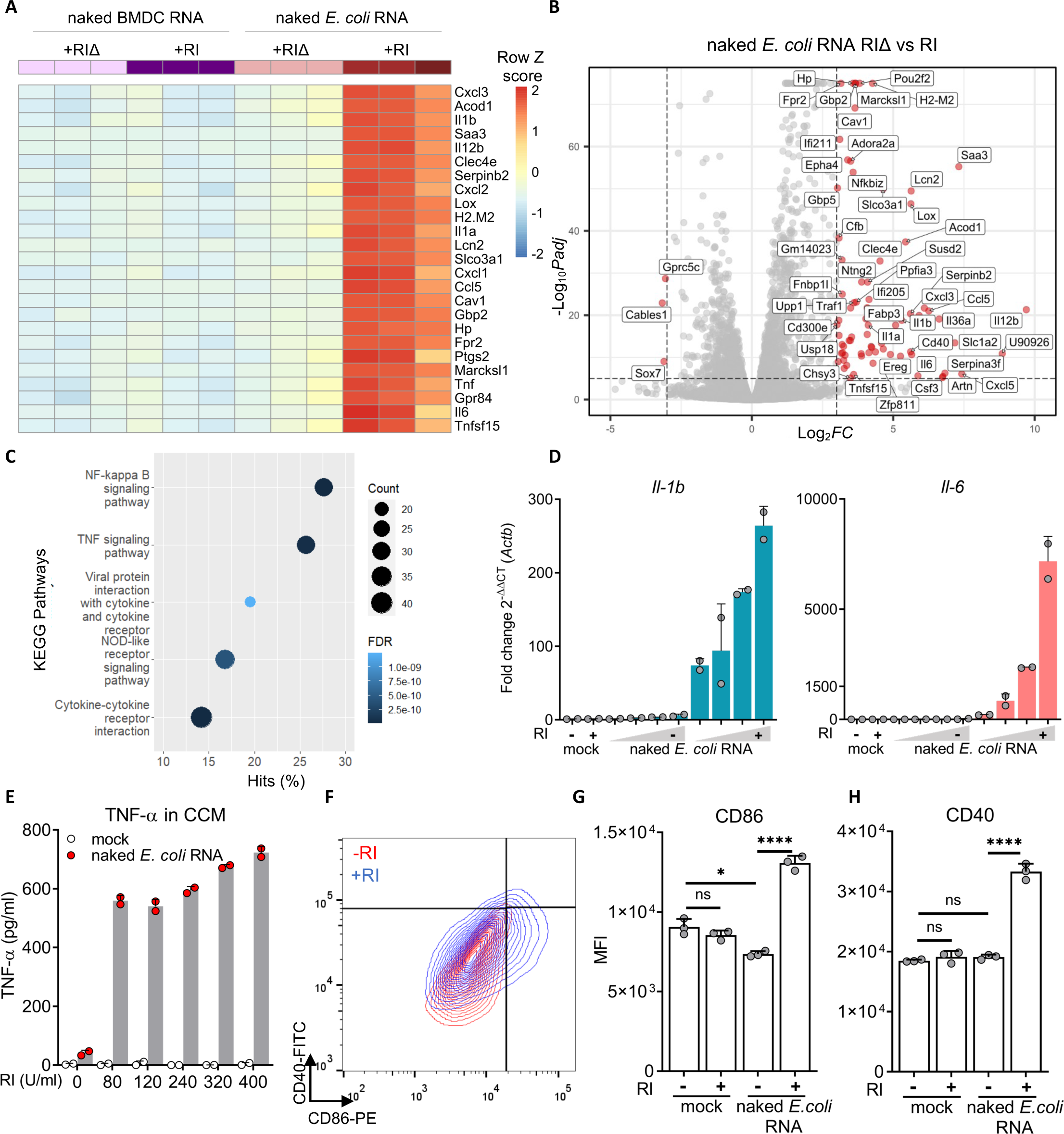
Naked bacterial exRNA activates BMDCs in a RI-dependent manner. **A**) Heatmap showing top 25 most differentially regulated genes in BMDCs stimulated for 6 h with 100 ng / mL naked RNA from *E. coli* or from BMDCs in the presence of RNase inhibitors (+RI) or a thermally-inactivated RI instead (+RIΔ). **B)** Volcano plot of differentially expressed genes (p-adj < 1×10^−3^, |log_2_FC| ≥ 3) between BMDCs stimulated with naked total *E.coli* RNA (+RI vs +RIΔ). **C)** Pathway enrichment analysis corresponding to BMDCs stimulated with naked *E. coli* RNA + RI. **D)** *Il-1b* and *il-6* expression measured by RT-qPCR in BMDCs stimulated with varying doses (1; 5; 10 or 25 μg / mL) of naked *E. coli* RNA, with or without RI. DPBS was used as negative control. **E)** TNF-α secreted by BMDCs after stimulation (6 h) with 1 μg / mL naked *E. coli* RNA (or DPBS) with varying doses of RI. **F-H)** Flow cytometry analysis of CD40 and CD86 in BMDCs stimulated for 24 h with 50 μg / mL naked *E. coli* RNA, with or without RI. Dot plots for treatments with naked total RNA **(F)** and median fluorescence intensity of CD86 **(G)** and CD40 **(H)** of all treatments are shown.

Previous results were further confirmed by RT-qPCR. All tested genes (*Cxcl10*, *Ifit2*, *Il1a*, *Il1b*, *Il6*, *Oas3*) showed a dose-dependent induction in the presence of RI (**Figure 1D** and **S3**). Additionally, pretreatment of naked *E. coli* RNA with RNAse1 completely abrogated the observed effects regardless of RI addition (**Figure S2B** and **S2C).** Interestingly, BMDCs exposed to naked *E. coli* RNA released high levels of TNF-α (**Figure 1E**) and increased the expression of cell surface markers associated with DC activation, such as CD40 and CD86 (**Figure 1F-1H**) only in the presence of RI. Overall, these results demonstrate that naked bacterial RNA can be specifically sensed by murine BMDCs, but this recognition cannot be observed if extracellular RNases are not previously inhibited.

### RI enhances the endocytic uptake of naked RNA by extracellular RNA stabilization

To understand whether naked exRNA sensing requires prior internalization, we incubated BMDCs with Dynasore, a dynamin inhibitor that blocks clathrin-dependent endocytosis and macropinocytosis (Preta et al. 2015), before the addition of naked exRNAs. Upregulation of *Il6* in response to bacterial exRNA was strongly inhibited in the presence of the drug (**Figure 2A**). Of note, whereas *Il6* induction was strongly dependent on RI (as previously observed; **Figure 1D**), endocytosis of Dextran-AF647 was independent of RI (**Figure 2B**). These results demonstrate that the main mechanism by which RI facilitates naked exRNA uptake is by stabilizing RNA in the extracellular space, and not by increasing endocytosis rates.

**Figure 2.**
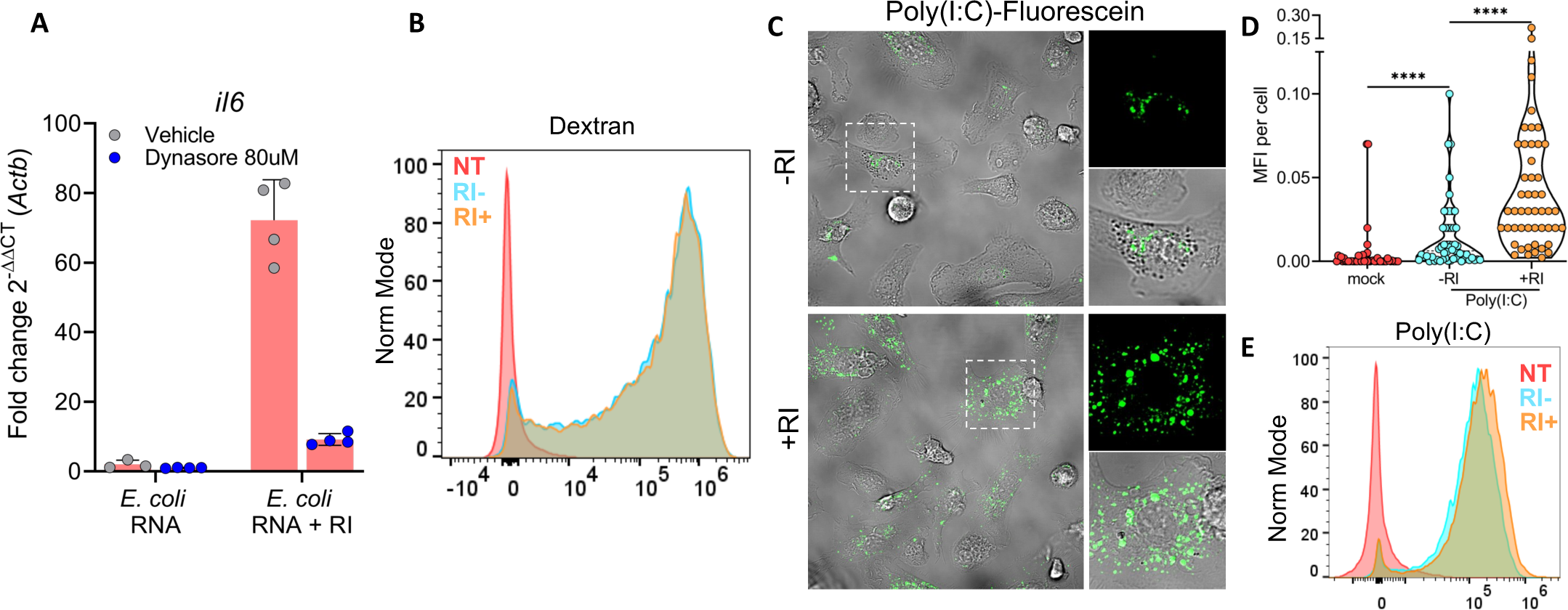
RI enhances the endocytic uptake of naked RNA by extracellular RNA stabilization. **A**) *il6* expression by RT-qPCR in BMDCs stimulated for 6 h with 1 μg / mL of naked *E. coli* RNA, with or without RI, in the presence of Dynasore (blue circles) or vehicle (grey circles). **B)** Flow cytometry analysis of BMDCs untreated or incubated with Dextran-AF647 for 1 h, with or without RI. **C,D,E)** Confocal microscopy **(C, D)** and flow cytometry analysis **(E)** of BMDCs cultured stimulated for 1h with 0.5 μg / mL naked poly(I:C)-Fluorescein.

To directly visualize internalized naked RNA molecules, BMDCs were stimulated with fluorescein-labeled RNAs, and imaged by confocal microscopy. To ensure detection in FBS-containing media, we used a synthetic double-stranded RNA analogue (poly(I:C)) due to its higher stability against ribonucleases. Interestingly, fluorescence signal appeared as discrete puncta, suggesting poly(I:C) localization in endosomal compartments. In agreement with previous results (**Figure 1**), a notable enhancement of this processes was observed upon inhibiting extracellular RNases (**Figure 2C** and **2D)**. This observation was orthogonally confirmed by flow cytometry (**Figure 2E** and **S4)**. Overall, these results position internalized naked RNAs, at least transiently, inside endosomes.

### A bacterial ribosomal RNA-derived TLR13 agonist activates RI-treated BMDCs

The localization of internalized exRNAs in endosomal compartments (**Figure 2**), and the potential role of the NF-κB pathway in BMDC response to naked exRNA (**Figure 1C**), suggests endosomal pattern recognition receptors (PRRs) are involved in naked exRNA sensing (**Figure 3A**).

**Figure 3.**
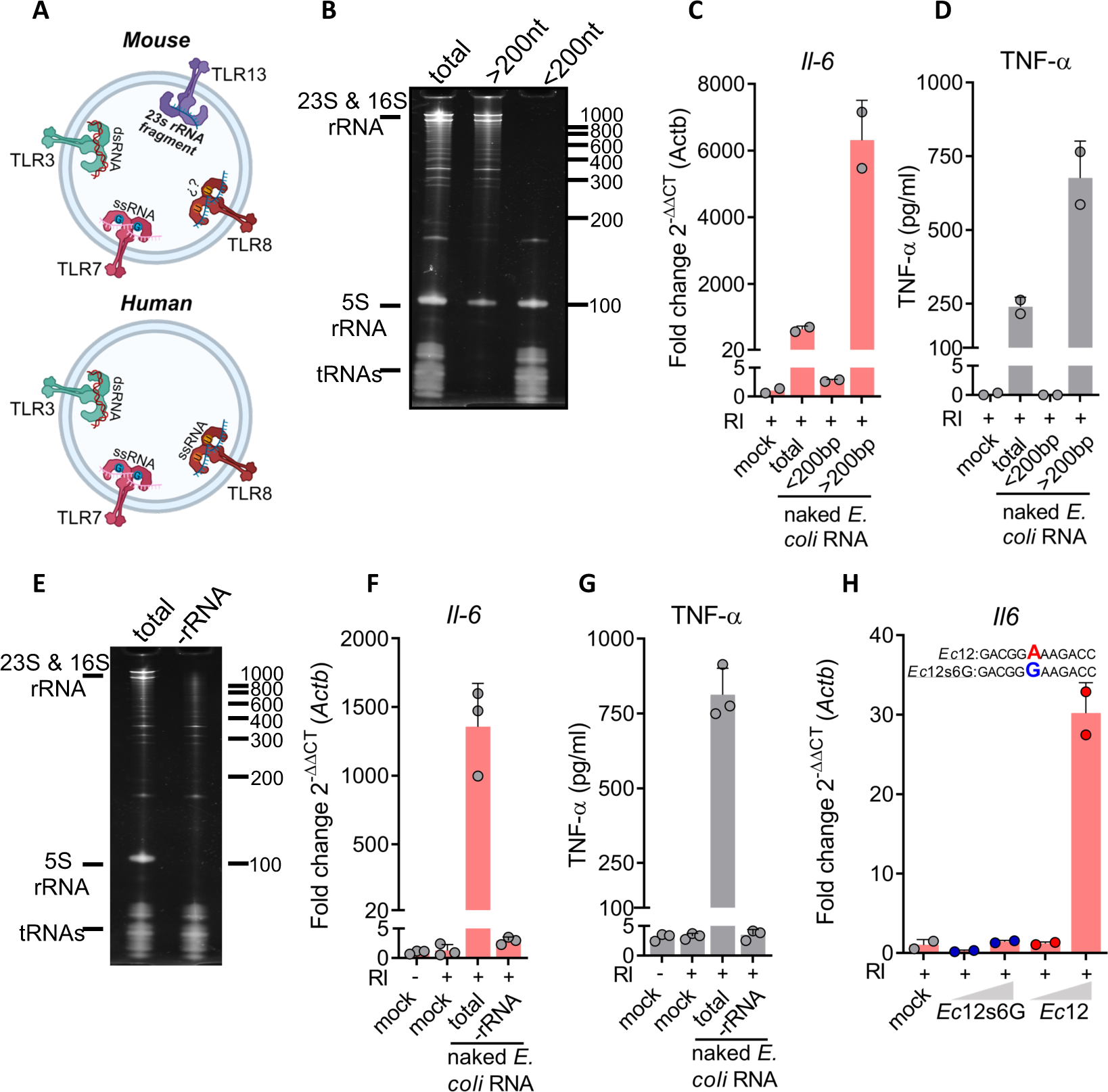
A bacterial ribosomal RNA-derived TLR13 agonist activates RI-treated BMDCs. **A**) Diagram of RNA sensing endosomal TLRs in mouse (top panel) and human (bottom panel). TLR3 sense double stranded RNA (dsRNA) larger than 40bp, TLR7 and TLR8 (in humans) sense short single stranded RNA (ssRNA) in the presence of Guanosine (G) or Uridine (U), respectively and TLR13 recognizes a ssRNA fragment derived from 23s bacterial rRNA **B)** Denaturing PAGE of *E. coli* total RNA (total) or fractioned into “small RNAs” of less than 200 nt (<200nt) or “large RNAs” of more than 200 nt (>200nt). **C,D)** *Il-6* expression (RT-qPCR) and levels of secreted TNF-α (ELISA) of BMDCs stimulated with 1 μg / mL of naked total *E. coli* RNA, “small RNAs” or “large RNAs”, all in the presence of RI. Mock: DPBS. **E)** Denaturing PAGE of total and ribosomal RNA-depleted (-rRNA) RNA from *E. coli*. **F,G)** *Il-6* expression (RT-qPCR) and levels of secreted TNF-α (ELISA) of BMDCs stimulated with 1 μg / mL of naked total or rRNA-depleted *E. coli* RNA, in the presence of RI. **H)** *Il6* expression (RT-qPCR) in BMDCs stimulated with 1 or 5 μg / mL of synthetic unmodified naked Ec12 RNA (a known agonist of TLR13), or its mutated version (Ec12s6G), both in the presence of RI. Mock: RPMI.

To infer which receptor-ligand interaction is mostly responsible for the observed effects, we separated total *E. coli* RNA into a small (< 200 nt) and a large (> 200 nt) fraction (**Figure 3B**) and observed that only naked RNAs longer than 200 nt triggered the production of proinflammatory cytokines at both the RNA and the protein levels (**Figure 3C** and **3D**). Considering that RNAs longer than 200 nt are mainly ribosomal RNAs (rRNAs), we speculated that extracellular bacterial rRNAs were triggering these effects. Indeed, probe-based selective depletion of bacterial rRNAs abrogated both *Il6* transcription and TNF-α release, despite using equal RNA concentrations across conditions (**Figure 3E-3G**). This strongly suggests the involvement of TLR13, a mouse-specific endosomal PRR that recognizes bacterial 23S ribosomal RNA and that is expressed in dendritic cells (Hidmark et al. 2012; Oldenburg et al. 2012).

A 23S rRNA segment, encompassing 12 nucleotides, was previously identified as the ligand of TLR13 in *S. aureus* (Oldenburg et al. 2012). We confirmed that this sequence, named here as *Ec12*, is conserved in *E. coli*. Interestingly, BMDCs stimulated with a synthetic and unmodified naked RNA comprising the sequence of *Ec12* triggered *Il6* expression, in the presence of RI (**Figure 3H).** Strikingly, this response was completely abrogated when adenosine in position six of *Ec12* was substituted for guanosine (*Ec12s6G)*, a substitution that is known to affect TLR13 recognition (Oldenburg et al. 2012). These results suggest the involvement of TLR13 in the recognition of endosomally internalized naked bacterial RNAs in murine BMDCs.

Overall, using naked *E. coli* RNA and BMDCs as a model, we have shown that extracellular RNAs can gain access to the endocytic pathway and trigger activation of endosomal RNA sensors when extracellular RNases are inhibited.

### Naked exRNA triggers MAVS-dependent cytosolic RNA sensors in human macrophages

To study whether other nonvesicular exRNAs could also activate BMDCs, we used the synthetic dsRNA analogue poly(I:C), a known agonist of TLR3 that can also engage the cytosolic RNA sensors RIG-I and MDA5 (Dauletbaev et al. 2015; McCartney et al. 2009) (**Figure 4A**), Interestingly, when BMDCs were incubated with naked extracellular poly(I:C), there was a strong increase in the expression of *Cxcl10* and *Ifit2* (**Figure 4B** and **4C**), two genes known to be induced by poly(I:C) in human DCs (**Figure S5A**). Unlike the *Cxcl10* and *Ifit2* response to naked bacterial exRNA (**Figure S3**), high poly(I:C) doses moderately stimulated BMDCs in the –RI condition. However, this response was markedly potentiated in the presence of RI (**Figure 4B** and **4C**).

**Figure 4.**
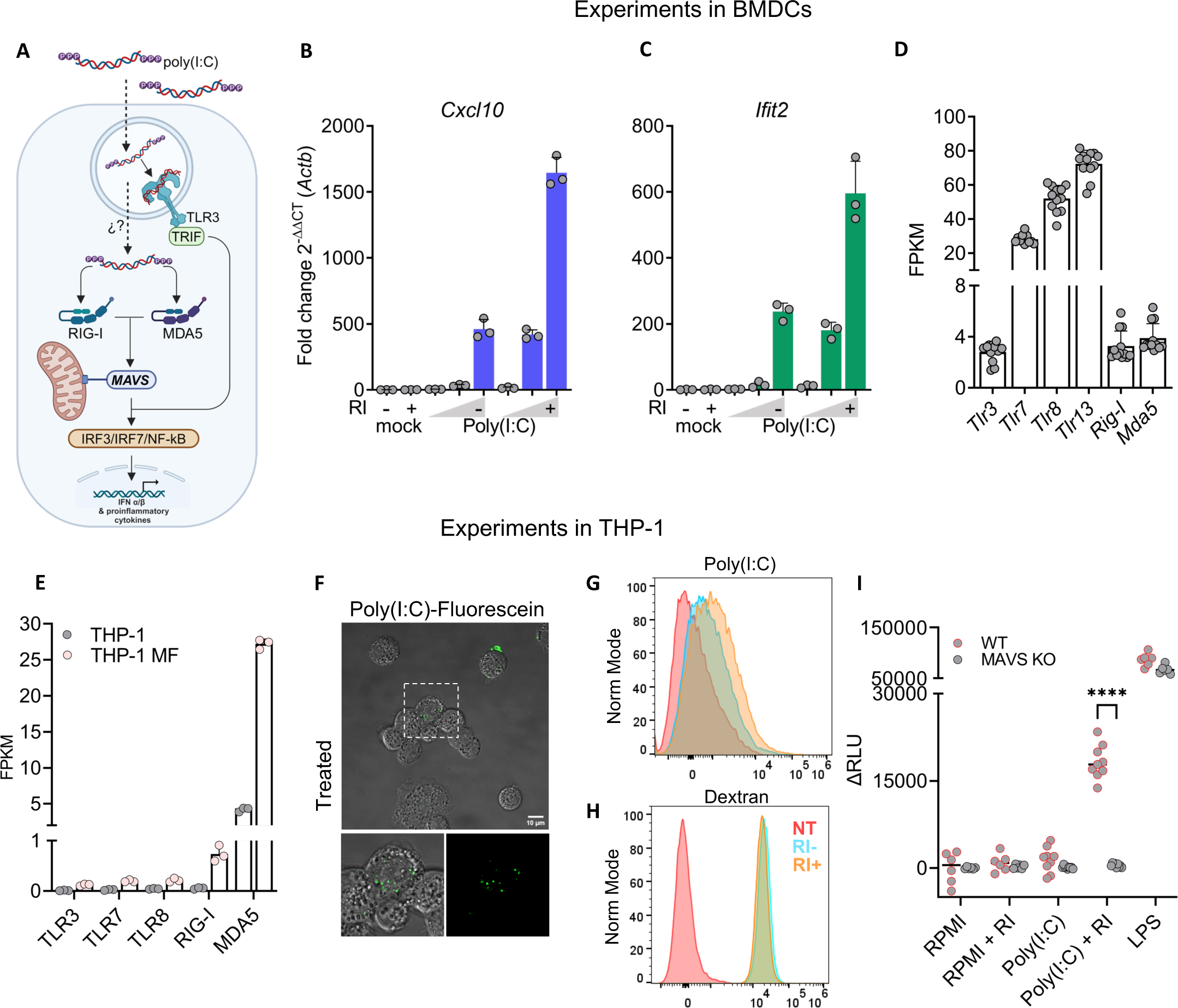
Naked exRNA triggers MAVS-dependent cytosolic RNA sensors in human macrophages. **A**) Diagram of RNA sensing endosomal and cytosolic receptor capable of recognizing naked poly(I:C) in BMDCs and THP-1 cells**. B,C)** BMDC expression of *Cxcl10* **(B)** and *Ifit2* **(C)** by RT-qPCR after 6 h stimulation with varying doses (0,1; 1; 10 μg/ml) of naked poly(I:C), with (+) or without (-) RI. RPMI was used as negative control. **D)** Expression (FPKM) of cytosolic and endosomal RNA pattern recognition receptors in BMDCs. **E)** Expression of cytosolic and endosomal RNA-specific pattern recognition receptors, in THP-1 cells or in THP-1-derived macrophages. Data was obtained from GEO: GSE130011. **F)** Confocal microscopy of THP-1 monocytes cultured in 10% FBS stimulated for 30 min with 0.5 μg / mL naked poly(I:C)-Fluorescein. **G, H)** Flow cytometry analysis of THP-1 monocytes cultured in 10% FBS for 2 h with naked poly(I:C)-Fluorescein (top panel) or with Dextran-AF647 (bottom panel), in the presence or absence of RI. **I)** Relative Luciferase activity corresponding to IRF pathway activation in reporter THP-1 Dual wild type or Mavs knockout cells after stimulation for 24 h with 5 μg / mL of naked poly(I:C), with or without RI. RPMI and 200 ng / mL LPS were used as negative and positive controls, respectively.

Although naked poly(I:C) appeared to be localized predominantly inside the endocytic compartment (**Figure 2C**), we also considered the possibility that at least a fraction of the internalized molecules could reach the cytosol, presumably by endosomal escape. Because BMDCs express both endosomal and cytosolic sensors capable of recognizing poly(I:C) (**Figure 4A** and **4D**), we instead used human THP-1 derived macrophages. Both the parental THP-1 cell line, as well as PMA-differentiated THP-1 macrophages, lack TLR3 but express high levels of the cytosolic RNA sensors RIG-I and MDA5 (**Figure 4E**). Hence, any response to poly(I:C) in these cells can be attributed, at least in theory, to cytosolic RNA sensors.

We first confirmed that THP-1 cells can internalize fluorescein-labeled naked poly(I:C) (**Figure 4F**) and that RI enhanced this process (**Figure 4G** and **S6**) without altering endocytosis rates in this cell line (**Figure 4H**). Then, we used a reporter THP-1 cell line that expresses a secreted luciferase under the control of IRF transcription factors. The IRF pathway is activated in response to type I interferons or several PRR agonists, including RIG-I and MDA5, which are both dependent on the mitochondrial antiviral-signaling protein (MAVS). Interestingly, naked extracellular poly(I:C) had no effect on reporter THP-1 macrophages. However, in the presence of RI, poly(I:C) triggered the IRF pathway in these cells (**Figure 4I**). Furthermore, this response was completely abrogated in MAVS knockout (KO) cells. This behavior cannot be attributed to a non-responsive KO cell line, as they behaved similarly to wild-type cells when using LPS, a MAVS-independent IRF pathway inductor.

Overall, these results demonstrates that naked exRNAs can also reach the cytosol, where the MAVS pathway is located, presumably by endosomal escape. In addition, we provide definitive proof of gymnotic RNA uptake with unmodified RNA molecules.

### RI facilitates protein translation after gymnotic uptake of mRNA

Our previous observations suggest that naked exRNA can, in the presence of RI, be internalized by cells and reach the cytosol after escape from the endosomal network. To assess the generalizability of this phenomenon, we studied gymnotic uptake of mRNAs in different human and murine cell types. Because translation occurs exclusively in the cytosol, observation of protein synthesis would provide conclusive evidence of gymnotic uptake, possibly involving endosomal RNA escape. We synthesized capped and polyadenylated *nanoLuc* mRNA by in vitro transcription (IVT) and confirmed its translatability using a rabbit reticulocyte extract (**Figure 5A**). Then, purified *nanoLuc* mRNA was added to murine BMDCs either with or without RI. Surprisingly, when extracellular RNases were inhibited, the intracellular levels of intact full-length mRNA molecules increased by 40-fold (**Figure 5B**). After 24 hours, we detected nanoLuc protein exclusively in BMDCs when extracellular RNases were inhibited (**Figure 5C**). To further underscore the influence of extracellular RNases on *nanoLuc* mRNA uptake and translation, we subjected BMDCs to varying concentrations of FBS and RI prior to their incubation with naked extracellular mRNA. As expected, increasing doses of RI stimulated translation, while increasing percentages of FBS had the opposite effect (**Figure 5D**). Beyond 10% FBS, RNase activity was sufficiently high to abolish *nanoLuc* translation, even in the presence of RI (**Figure 5D**).

**Figure 5.**
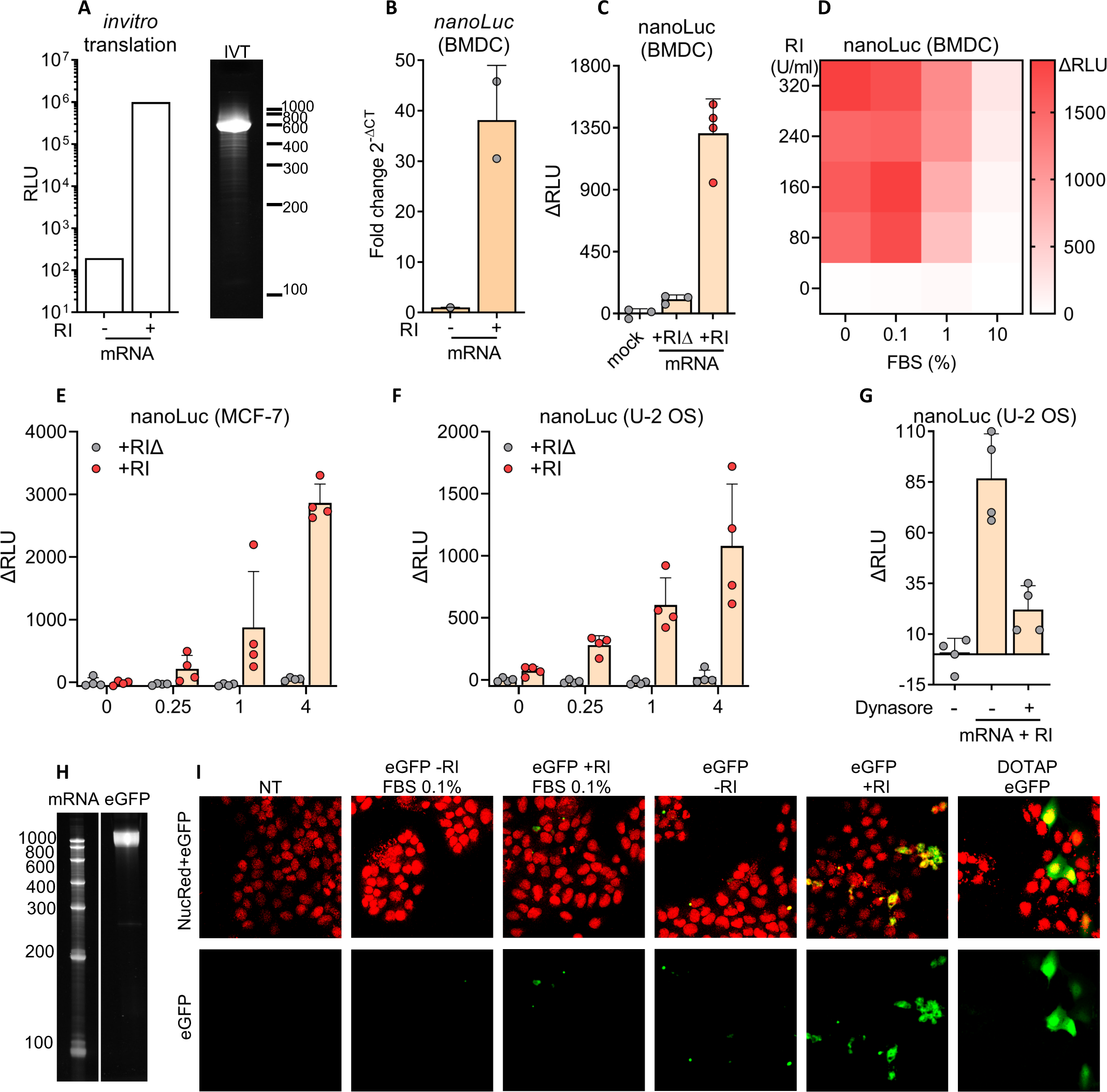
RI facilitates protein translation after gymnotic uptake of mRNA. **A**) Bioluminescence of in vitro translated nanoLuc protein (left) and denaturing PAGE of in vitro-transcribed nanoLuc mRNA (right). B) BMDCs cultured without FBS were incubated with 1 μg / mL naked nanoLuc mRNA with or without 80 U / mL RI. After 6 h, intact intracellular nanoLuc mRNA was quantified by RT-qPCR. **C)** After 24 hs, nanoLuc protein levels were quantified by bioluminescence. **D)** Heatmap showing nanoLuc intracellular protein levels detected by bioluminescence in BMDCs (cultured with varying doses of FBS) as a function of RI concentration. **E,F)** MCF-7 **(E)** and U-2 OS **(F)** cells cultured in 0.1% FBS were incubated with increasing doses of nanoLuc mRNA (0; 0,25; 1; 4 μg / mL) with 80 U / mL RI. After 24 hs, NanoLuc protein levels were quantified by bioluminescence. **G)** U-2 OS cells cultured in 0.1% FBS were incubated with nanoLuc mRNA and RI in the presence of 80 μM Dynasore or vehicle. After 24 hs, nanoLuc protein levels were determined by bioluminescence. **H)** Denaturing PAGE corresponding to in vitro-transcribed eGFP mRNA. **I)** Confocal microscopy of U-2 OS cells incubated with 5 μg / mL naked eGFP mRNA with or without 80 U / mL RI at the indicated FBS dose. Non-treated cells and DOTAP-transfected cells were used as negative and positive controls, respectively. Green channel: eGFP, Red channel: Nuclei.

Since BMDCs are professional phagocytic cells, we sought to determine if uptake occurred for other cell types. Interestingly, several human epithelial cell lines could uptake and translate nanoLuc mRNA in the presence of RI (**Figure 5E** and **5F**). As previously observed in murine BMDCs, nanoLuc translation in human epithelial cells was abrogated in the presence of endocytosis inhibitors (**Figure 5G**). Additionally, using confocal microscopy, we could detect that U-2 OS cells are also able to capture and translate naked eGFP mRNA, but only if cells are cultured in the absence of FBS and RI is added to the media (**Figure 5H** and **5I**).

### Compartment-specific RNase activity modulates naked exRNA-induced inflammation

We have observed that naked exRNA induces pro-inflammatory responses in cultures of human and murine immune cells, and that these responses are modulated by extracellular RNase activity. To assess whether naked exRNAs could also induce inflammatory responses in vivo, we injected purified *E.coli* RNA into the peritoneal cavity of immunocompetent mice (Figure 6A). Interestingly, naked bacterial RNA induced the disappearance of resident large peritoneal macrophages and the recruitment of circulating monocytes at 24 h; a hallmark of peritoneal inflammation (Louwe et al. 2021) (**Figure 6B** and **S7A**). Of note, in contrast with our previous in vitro results, the addition of RI had negligible effects (**Figure 6C**, and **S7B**). To understand this discrepancy, we assessed the RNase activity of the peritoneal cavity compared with an identical dilution of FBS (3.3%), assuming a peritoneal volume of 100 μl (Hartveit and Thunold 1966) that was diluted to 3.1 mL by addition of peritoneal wash buffer. Notably, RNA decay assays showed that FBS contains a high RNase activity even at such a high dilution rate, with a peritoneal wash supporting almost 30-fold slower decay (**Figure 7A**). Thus, the fact that naked bacterial RNA is proinflammatory in the peritoneal cavity irrespective of RI addition suggests that, in RNase poor environments, naked extracellular RNA could be directly sensed by the immune system. Following this line of reasoning, these results imply that extracellular RNases may have evolved to degrade potentially life-threatening systemic inflammatory responses induced by naked exRNAs.

**Figure 6.**
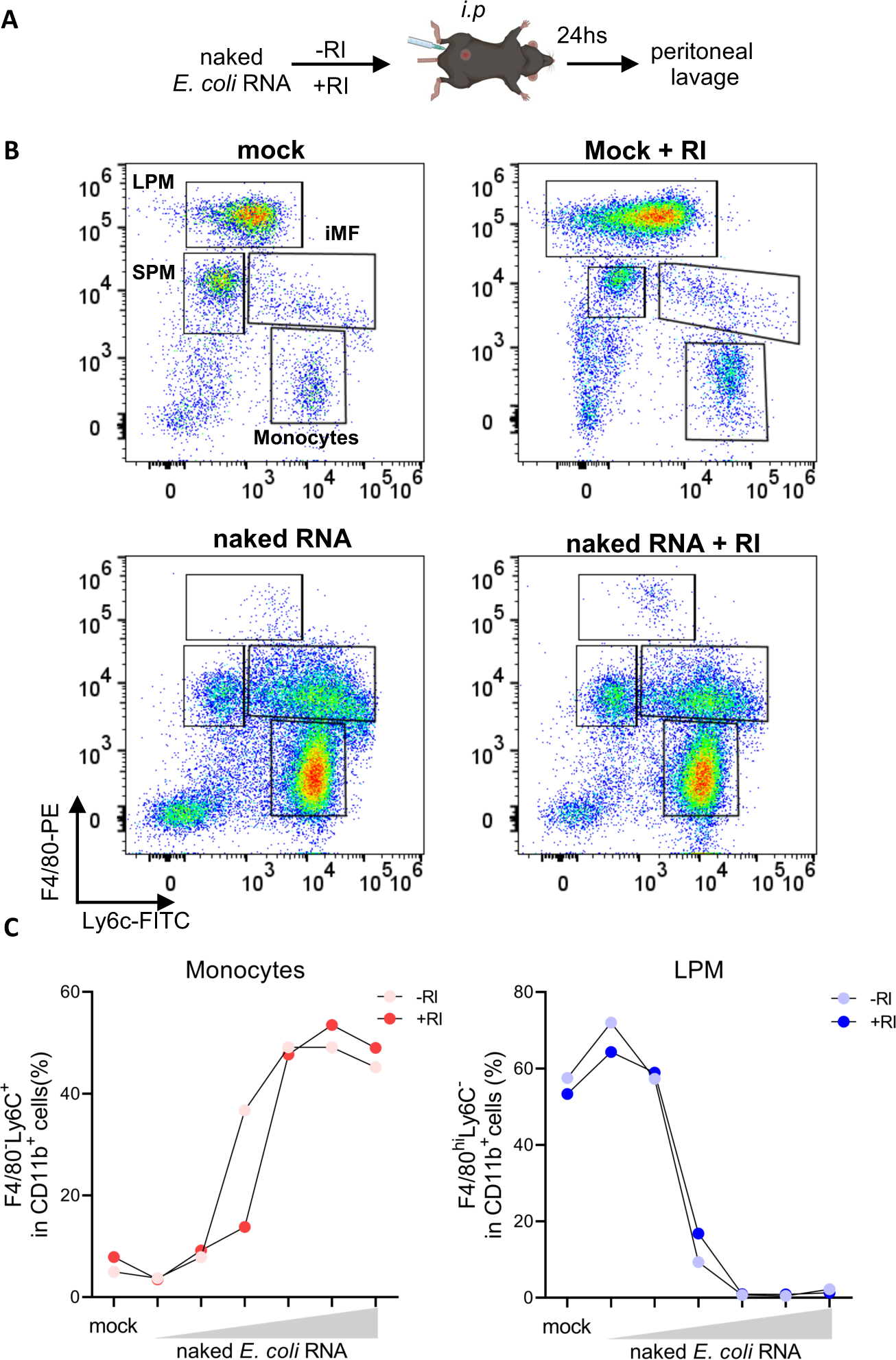
Naked exRNA is intrinsically bioactive in the peritoneal cavity. **A**) Experimental scheme. **B)** Flow cytometry dot plots of F4/80 and Ly6c (gated in lineage-CD11b^+^) showing immune cell populations at the peritoneal cavity 24 h after intraperitoneal (i.p.) injection of PBS, PBS + RI, 25 μg of naked total *E. coli* RNA, or 25 μg of naked total *E. coli* RNA + 40 U RI. Resident large (LPM) and small (SPM) peritoneal macrophages, infiltrating monocytes (Mo) and inflammatory macrophages (iMF) are shown. Gating strategy is shown in **Figure S7**. **C)** Graphs showing the percentage of peritoneal infiltrating monocytes defined as Lin^−^CD11b^+^F4/80^−^Ly6c^+^ (left panel) and resident large peritoneal macrophages (LPM) defined as Lin^−^CD11b^+^F40/80^hi^Ly6c^−^ (right panel), 24 hs after *i.p* injections of DPBS (mock) or increasing doses of naked total *E. coli* RNA (0.025; 0.250; 1.25; 12.5; 25; 62.5 μg) in the absence or presence of RI.

**Figure 7.**
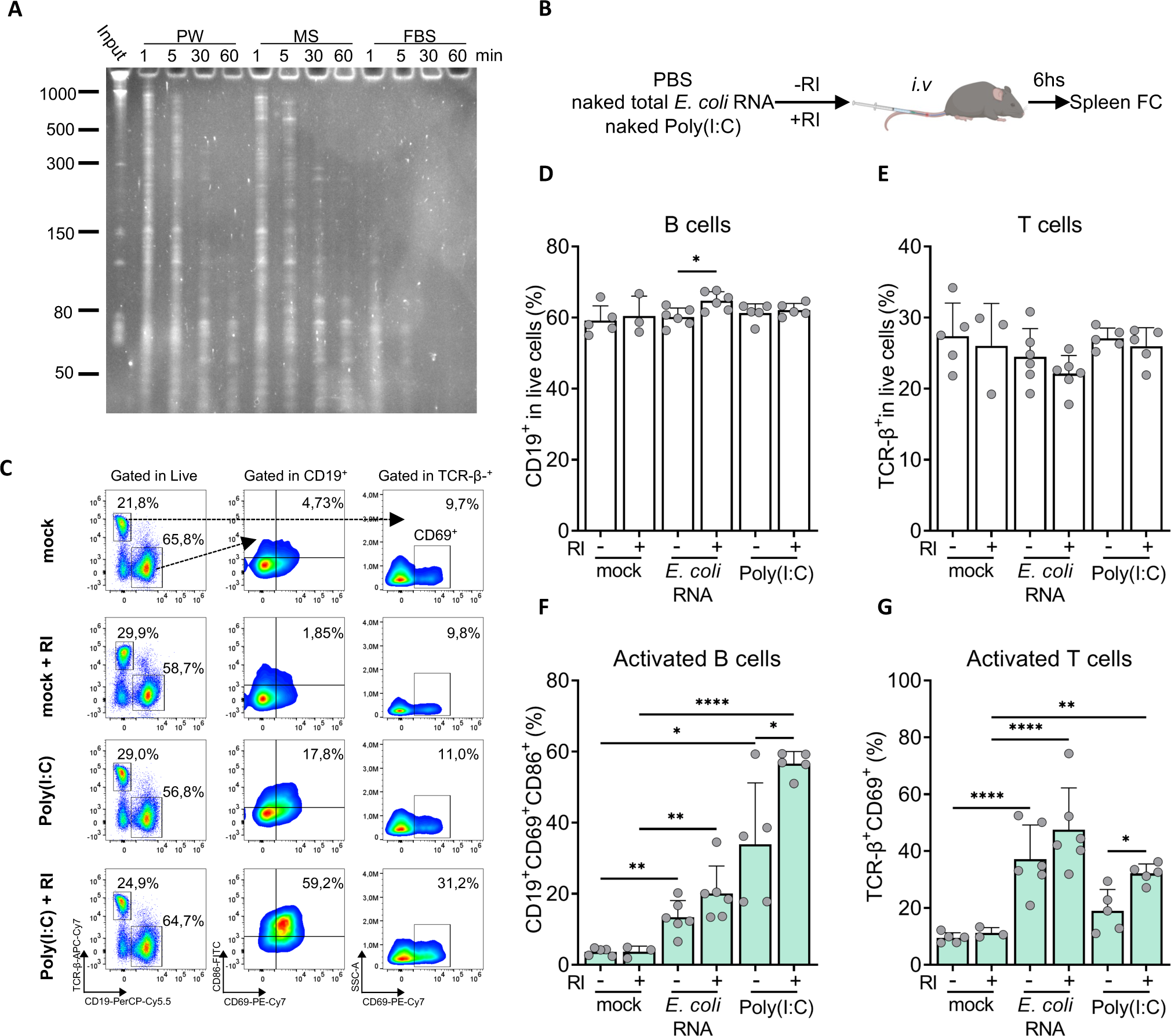
RI enhances the immunostimulatory effects of naked exRNAs circulating in the blood. **A**) RNA stability assays in different biofluids. U-2 OS RNA was incubated ex vivo in a peritoneal wash (PW), mouse serum (MS) or fetal bovine serum (FBS), and RNA degradation was analyzed after 1, 5, 30 or 60 minutes. All biofluids were diluted to 3.33 % (v/v) in PBS. **B)** Experimental scheme. **C-G)** Flow cytometry analysis of spleen cells 6 h after intravenous administration of either naked poly(I:C) (1 μg) or naked total *E. coli* RNA (10 μg), with or without 480 units of RI. Mock: PBS. Representative dot plots are shown in **(C)**. The percentage of total B cells (CD19^+^) and T cells (TCR-β^+^), gated in live cells, is shown in panels **D** and **E**. The percentage of activated B cells (defined as CD69^+^CD86^+^) and T cells (defined as CD69^+^), gated under their corresponding cell type, is shown in panels **F** and **G**.

To test the impact of circulating RNases in the regulation of systemic inflammation induced by exRNA, we intravenously injected both naked *E.coli* RNA and poly(I:C) (**Figure 7B**) with or without 480 units of RI; a dose sufficient to inhibit mouse serum RNases (**Figure S8**). Surprisingly, both naked exRNAs increased the percentage of activated (CD86^+^MHCII^+^) cDCs, pDCs, and macrophages in the spleen (**Figure S9**), as well as the percentage of activated splenic B cells (CD86^+^CD69^+^) and T cells (CD69^+^) (**Figure 7C** and **S10**), with minor or no changes in total cell numbers (**Figures 7D** and **7E**). Importantly, despite the lower RNase activity observed in mouse serum compared with FBS (**Figure 7A**), we could still recapitulate the effect of RI addition in the case of poly(I:C). In agreement with in vitro results, an increased population of activated B cells, T cells, macrophages, and pDCs was observed compared with poly(I:C) alone (**Figure 7F** and **7G**, and **S9B)**.

In summary, naked exRNA is intrinsically bioactive in RNase-poor environments such as the peritoneal cavity. In addition, these results suggest a key role for bloodborne extracellular RNases in the regulation of exRNA-induced systemic inflammation.

## DISCUSSION

RNA-mediated intercellular communication has been almost exclusively studied in the context of extracellular vesicles (EVs) (O’Brien et al. 2020), despite most exRNAs being present outside EVs in human biofluids (Arroyo et al. 2011; Turchinovich et al. 2011), cell-conditioned media (Tosar et al. 2015, 2020; Sork et al. 2021), and even in plants (Borniego and Innes 2023; Zand Karimi et al. 2022). A bias against the study of nonvesicular exRNAs can be explained by the widespread assumption that exRNA is unstable unless protected inside EVs. Additionally, EVs provide a mechanism for functional RNA delivery into recipient cells (O’Brien et al. 2020). In contrast, naked RNA is assumed to be incapable of penetrating the “billion-year-old barrier” comprising the plasma membrane and the membrane of endocytic vesicles (Dowdy 2017). By being refractory to the uptake of extracellular RNAs, cells could protect themselves from selfish genetic elements such as viroids and positive-stranded RNA viruses, while preserving their transcriptional identity.

However, several RNA species are stable in the extracellular space even when not associated with EVs. These resilient extracellular RNAs include ribonucleoprotein particles (RNPs) such as Ago2/miRNA complexes (Turchinovich et al. 2011; Arroyo et al. 2011; Geekiyanage et al. 2020), and protein-protected RNA fragments (LaPlante et al. 2023) derived from the ribosome (Tosar et al. 2020, 2022; Costa et al. 2023), U2 snRNPs (Tosar et al. 2022), and possibly Xist RNPs (Dou et al. 2024). Additionally, a population of intrinsically stable nicked tRNAs circulates in human biofluids, probably unprotected or naked (Costa et al. 2023).

Rather than being inert, naked extracellular RNAs can enter cells by gymnosis. It should be noted that this term was not defined in the context of a precise molecular mechanism (Stein et al. 2010); the name axiomatically refers functional naked RNA delivery into the cytosol (Zhang et al. 2018). Since best results in the original publication were obtained with locked nucleic acids (LNA)-containing phosphorothioate ASOs (Stein et al. 2010), gymnotic uptake was later considered as an emerging property of phosphorothioate-containing oligonucleotides (Dowdy 2017). Unmodified and longer naked RNAs are thought to lack any biological activity.

In this work, we provide a different interpretation: any RNA can enter cells by endocytosis, and even reach the cytosol, as long as its stability in the extracellular space is sufficiently high. The main factor limiting the concentration of naked extracellular RNAs are extracellular RNases, which are highly abundant and active in biofluids such as the human blood (Lomax et al. 2017; Costa et al. 2023). Phosphorothioate-contaning ASOs are resistant to degradation, and this may be why gymnotic uptake can occur for ASO-based biotherapeutics (Levin 2019). By adding a broad-range RNase inhibitor to cell culture media, we demonstrated that uptake can be extended to naked bacterial rRNAs, synthetic oligonucleotides, poly(I:C), and in vitro-transcribed mRNAs. This was shown in a variety of different cell types, including primary cultures of professional phagocytes and human epithelial cell lines.

We have shown that naked extracellular RNAs, when stabilized by RI, are intrinsically bioactive in recipient cells. They can engage with endosomal RNA sensors such as TLR13 (in BMDCs), but they can also escape from endosomes into the cytosol, where they can be recognized by MAVS-dependent cytosolic RNA sensors (at least in human THP-1 macrophages). More strikingly, two different mRNAs encoding reporter proteins could be internalized by gymnosis and still be capable of serving as templates for protein synthesis in both BMDCs and human epithelial cell lines. This suggests their capacity to perform endosomal escape because ribosomes are present only in the cytosol, including on the cytosolic face of membranous organelles such as the ER and mitochondria. It is important to mention that these mRNA translation assays are highly sensitive to extracellular RNases, because a single cleavage event would render an mRNA untranslatable. Hence, the effect of RI was more pronounced in these assays, and was potentiated by FBS depletion.

RNA endosomal escape in the absence of LNPs means that the “billion-year-old barrier” (Dowdy 2017) may actually be leaky under certain circumstances. Interestingly, while efficient translation of our GFP-encoding mRNA required the use of modified nucleotides to avoid recognition by TLR7 (Karikó et al. 2005), nanoLuc mRNA was translated even with unmodified uridine. These results encourage further basic research in naked mRNA therapies and vaccines. Although positive results were obtained after localized naked mRNA injections in organs with presumably low extracellular RNase activity (Wolff et al. 1990; Kreiter et al. 2010; Diken et al. 2011), co-administration of RI could enable more systemic delivery routes.

If pathogen-derived naked extracellular RNAs are potent pro-inflammatory molecules, how are potentially life-threatening systemic inflammatory responses avoided? For example, the complement system could induce bacterial lysis in the bloodstream, releasing high loads of pro-inflammatory bacterial rRNAs that could activate circulating leukocytes. This study strongly suggests that extracellular RNases are highly abundant in the bloodstream to avoid RNA-induced systemic inflammation. We showed this by demonstrating that RI co-administration enhances immune cell activation in the spleen after intravenous injection of naked RNA. Interestingly, RI was dispensable when a similar assay was performed in the peritoneal cavity, an important site of immune surveillance with much lower RNase activity. Thus, naked extracellular RNAs might convey important information for immune cells residing in tissues with relatively low blood irrigation and hence with low extracellular RNase content.

The immune system has multiple regulatory layers to ensure an optimal response to the different stimuli that immune cells could encounter. One key challenge is to differentiate self from non-self nucleic acids, which is achieved by regulating both the affinity (Brown et al. 2022) and the subcellular localization of endosomal TLRs (Lind et al. 2022; Mishra et al. 2024). Indeed, we observed a dramatically different response when comparing bacterial and murine RNAs incubated at the same concentration, both in the presence of RI. Surprisingly, the effect of RI itself was as dramatical as the capacity of BMDC RNA sensors (predominantly TLR13) to discriminate self from non-self RNAs. Thus, RNase-mediated control of exRNA stability lies on top of a series of layers comprising control of endocytosis, TLR loading and recognition, endosomal escape, and engagement with cytosolic RNA sensors.

Supporting the role of extracellular RNases in the regulation of inflammatory responses, several reports have shown that ectopic administration of extracellular RNases reduces inflammation in mouse models of Lupus-like disease (Sun et al. 2013), traumatic brain injury (Krämer et al. 2022), and myocardial infarction (Cabrera-Fuentes et al. 2014). In humans, sterile inflammation causing cell death and the release of RNPs into the extracellular space seems to be an important trigger of Systemic Lupus Erythematosus (Dou et al. 2024). In addition, polytrauma induces exRNA release followed by TLR7-dependent systemic inflammation in mice (Suen et al. 2023). The involvement of extracellular RNases in the onset and progression of autoimmune disease should therefore be investigated. If exRNA could be considered a damage-associated molecular pattern (DAMP), extracellular RNases might be also reasoned as a regulatory arm of the innate immune system.

On a more technical note, we would like to emphasize that intents to understand extracellular RNA biology using cell culture-based assays will inevitably fail if fetal or calf serum is used without RI supplementation. This technical problem has led to the wrong assumption that exRNA biology is relevant only in the context of EVs, or that the study of intracellular nucleic acid sensors require the use of transfection reagents (Sha et al. 2014). We envision that this study will renovate interest in nonvesicular extracellular RNAs in intercellular communication, particularly in the context of inflammatory and autoimmune diseases.

## CONCLUSION

This study clearly demonstrates that naked exRNA is intrinsically bioactive both in vitro and in vivo. Moreover, in the absence of RNases, exRNA can spontaneously enter cells and even escape from endosomal membranes, engaging both endosomal and cytosolic RNA sensors. Earlier studies may have failed to observe these phenomena due to the confounding effect of extracellular RNases. Furthermore, this study suggests that the multiple RNases that exist in the extracellular space may have evolved to avoid potentially harmful systemic inflammatory responses triggered by naked or nonvesicular exRNAs.

## ACKNOWLEDGMENTS

J.P.T., M.S., M.H. and A.C. are members of the Sistema Nacional de Investigadores (SNI) and Programa de Desarrollo de las Ciencias Básicas (PEDECIBA, MEC-UdelaR). The authors gratefully acknowledge the following core facilities and technological platforms at the Institut Pasteur of Montevideo for their support & assistance in the present work: Advanced Bioimaging Unit (UBA), Animal Biotechnology Unit (UBAL) and the Cell Biology Unit (UBC) with a special thanks to Karen Perelmuter and Paula Céspedes. The authors want to thank Sofía Russo, Daniela Olivera, Germán Galliussi and Valentina Perez for helpful suggestions, experimental help and insightful scientific discussions.

## METHODS

### RESOURCE AVAILABILITY

#### Lead contact

Further information and requests for resources and reagents should be directed to and will be fulfilled by the corresponding author, Juan Pablo Tosar.

## EXPERIMENTAL MODEL

### Animals

Male and female C57BL/6 mice, aged between 8 to 12 weeks, were used (Jackson Lab; Bar Harbor, ME). Mice were bred for up to 20 generations in a pathogen-free environment at the Laboratory Animal Biotechnology Unit of the Pasteur Institute of Montevideo. All experiments were performed under strict adherence to the guidelines set by the National Commission for Animal Experimentation. Experimental procedures were approved by the Ethics Committee on Laboratory Animals of the Pasteur Institute of Montevideo (protocol: 022-22). For primary cell cultures both male and female C57BL/6 mice were used. For in vivo experiments, C57BL/6 female mice were used.

### Cell lines

U-2 OS and MFC-7 cells were cultured in DMEM supplemented with 10% FBS without antibiotics, THP-1 cells were cultured in RPMI with glutaMAX supplemented with 10% FBS without antibiotics. THP-1-Dual and THP-1 Dual Mavs knockout reporter cell lines (Invivogen, USA) were cultured in RPMI with glutaMAX, supplemented with 10% FBS, 25 mM HEPES, 2mM Glutamine, 100 U / mL Penicillin-Streptomycin and 100 μg / mL Normocin. Selection pressure was maintained with 10 μg / mL Blasticidin and 100 μg / mL Zeocin. THP-1, THP-1-Dual and THP-1 Dual Mavs knockout monocytes were differentiated to macrophages by 3 h stimulation with 40 nM Phorbol 12-myristate 13-acetate (PMA), followed by 48-72 h incubation without PMA.

### Primary cell culture

Differentiated Bone Marrow Derived Cells, containing dendritic cells (BMDCs), were cultured from bone marrow cell precursors as described in (Segovia et al. 2019). Briefly, bone marrow cells were collected form 8–12 weeks-old C57Bl/6 mice and differentiated with 0.4 ng / mL GM-CSF for 8 days in RPMI supplemented with 10 % heat-inactivated FBS, 2mM glutamine, non-essential amino acids, 1 mM sodium pyruvate, 10 mM HEPES, 0.05 mM β*-*mercaptoethanol and 100 U / mL Penicillin-Streptomycin. At day 8, cells were harvested and used as needed.

## METHOD DETAILS

### Total E. coli RNA purification and fractionation

Total RNA was extracted from exponentially growing *E. coli* DH5α cells with OD = 0.6. Cells were lysed in RLT buffer supplemented with 20 mM DTT using a bullet blender homogenizer (Next Advance). The homogenate was precipitated with 2.5 volumes of 100% acetone. The resulting pellet was resuspended in TRIzol, and total RNA was purified from the aqueous phase using a Monarch RNA cleanup kit. To isolate large (> 200 nt) and small (< 200 nt) RNAs, total RNA was sequentially bound to Monarch spin columns using binding buffer with 33% ethanol (allows binding of only >200 bp RNAs) and 66% ethanol (binds all RNAs). Additionally, 23S, 16S and 5S ribosomal RNAs were selectively removed with a NEBNext rRNA Depletion Kit for bacteria. When required, total *E. coli* RNA was degraded with recombinant human RNase 1.

### RNA-seq and bioinformatic analysis

BMDCs (300,000 cells per well) cultured in growth medium with 10% FBS and supplemented with 10 µg / mL Polymyxin B were stimulated with 100 ng / mL naked total RNA from *E. coli* or naked total self-RNA from BMDCs (isolated from cultures at day 8), in the presence of either 80 U / mL RI or heat-inactivated RI (RIΔ) for 6 h. Afterwards, cells were lyzed with TRIzol and RNA present in the aqueous phase after addition of chloroform was purified with a Monarch RNA cleanup kit. RNA integrity was confirmed by electrophoresis in 10% TBE-urea polyacrylamide gels. RNA was precipitated with 0.1 volumes of 3M sodium acetate pH=3.2 and 2 volumes of 100% ethanol and shipped to Macrogen (Korea) for paired-end mRNA sequencing (TruSeq Stranded mRNA, Illumina). Then, FastQ files containing paired-end sequencing information were mapped to the mouse genome (GRCm39) using STAR (Dobin et al. 2013) (default settings). Mapping quality was assessed with QualiMap (Okonechnikov et al. 2016) (default settings). Expression of mRNAs across samples was analyzed with FeatureCounts (Liao et al. 2014) (default settings and paired-end reads counted as a single fragment) using Ensembl mouse annotation (Release v109). Differential expression across samples was computed with DESeq2 (Love et al. 2014) (default settings). Heatmaps showing topmost differentially expressed genes were calculated using rlog-transformed counts and the pheatmap R package and row-scaled. Volcano plots and MA-plots were constructed using EnhancedVolcano and ggpubr R packages, respectively. Differentially expressed genes between conditions of naked *E. coli* RNA with RI and naked *E. coli* RNA with inactivated RI having fold change > 2, adjusted p-value < 1×10^−5^ and base mean > 16 were used for pathway enrichment analysis using ShinyGO (v0.77) (Ge et al. 2020). To determine expression differences between RNA sensors, fragments per kilobase per million (FPKM) were calculated for *Tlr3*, *Tlr7, Tlr8, Tlr13, Rig-I, Mda-5*.

### Bioinformatic analysis of publicly available datasets

Publicly available transcriptomic datasets of THP-1 cells differentiated to THP-1 macrophages with PMA (GEO: GSE130011) (Green et al. 2020) and human monocyte-derived dendritic cells stimulated with different TLR agonists (GEO: GSE125817) (Johnson et al. 2020) were downloaded from the NCBI SRA repository and subjected to a similar bioinformatic pipeline as previously described. In these cases, reads were mapped to the human genome (hg38) and expression of mRNAs across samples was analyzed with FeatureCounts using Ensembl human annotation.

### Cell stimulation

BMDCs seeded in 24 well plates (300,000 cells/well) cultured with growth medium containing 10% FBS were stimulated for either 6 or 24 h with the following RNAs at the indicated dose: total *E coli* RNA, rRNA-depleted *E coli* RNA, size-fractionated *E. coli* RNA, synthetic RNAs Ec12 and Ec12s6G (sequences provided in the key recourses table) and high molecular weight poly(I:C), together with 80 U / mL RI, thermally inactivated RI (RIΔ), or without RI. RPMI or DPBS were used as negative controls and 100 ng / mL R848 was used as a positive control. Whenever bacterial RNA was used, cells were cultured with 10 µg / mL Polymyxin B to quench any endotoxins remaining after RNA purification. After 6 h, the expression of *Il1b*, *Il1a*, *Il6, Cxcl10, Ifit2,* and *Oas3* was analyzed by RT-qPCR (See primers in **Table S1**). Secreted TNF-α was quantified in cell-conditioned medium by ELISA. After 24 h, cell surface expression of CD40 and CD86 was analyzed by flow cytometry. To study involvement of dynamin-dependent endocytosis on internalization and responses of naked inflammatory RNAs, BMDCs were stimulated for 6 h with 1 μg / mL naked total RNA with or without 80 U / mL RI in the presence of 80μM Dynasore or vehicle (DMSO). In other experiments, BMDCs were stimulated with 0.5 µg / mL naked Fluorescein-labeled poly(I:C) or Dextran-AF647 in the presence or absence of 120 U / mL RI for 1 h, and analyzed by confocal microscopy and flow cytometry. THP-1 dual wild type and *Mavs* knockout cells (Invivogen) seeded in 96 well adherent plates and cultured with growth medium containing 10% FBS were differentiated to macrophages with PMA. Then THP-1 derived macrophages were stimulated for 24 h with 5 μg / mL naked poly(I:C), with or without 80U / mL RI. RPMI and 200 ng / mL LPS were used as negative and positive controls, respectively. After 24 h, luciferase activity in the medium was detected following manufacturer’s instructions using a LUMIstar Optima luminometer. In other experiments, THP-1 cells were stimulated with 0.5 µg / mL naked Fluorescein-labeled poly(I:C) or Dextran-AF647 in the presence or absence of 120 U / mL RI for 30 or 120 min and analyzed by confocal microscopy and flow cytometry.

### RT-qPCR

BMDC RNA was extracted with TRIzol following manufacturer’s instructions and quantified spectrophotometrically. cDNA was synthesized from 300-1000 ng total RNA using M-MLV reverse transcriptase (Thermo). Briefly, total RNA was treated with 2 units of DNase I for 20 min at 25°C to eliminate genomic DNA, in a reaction volume of 10 μl. DNase I was inactivated by addition of EDTA at a final concentration of 4.5 mM and heating at 75°C for 10 min. Then, RNA was reverse-transcribed in a 20-μl reaction volume using oligo(dT)_18_ primers, following manufacturer’s instructions and employing a two-step PCR program: 52 min at 37°C, followed by 15 min at 70°C. Then, cDNA was diluted 1/5 and stored at –20°C until use. Gene expression was quantified by qPCR in a QuantStudio 3 Real-Time PCR System (Applied Biosystems) employing a FastStart Universal SYBR Green Master Mix. Briefly, a 10-μl reaction was carried out using 2 μl of cDNA, a primer mix at 0.3 μM final concentration, and 5 μl of a SYBR Green Master Mix. The qPCR program was: 2 min at 50°C, followed by 10 min at 95°C, followed by 40 cycles of 15 s at 95°C and 1 min annealing/extension at 60-65°C, and a final melt curve stage. Comparative gene expression analysis was calculated using the 2^−ΔΔCT^ method. Data was normalized against mock treatment (RPMI or DPBS) and *actb* was used as a housekeeping, reference gene.

### In vitro transcription

mRNAs coding for *nanoLuc* or *eGFP* were synthesized by in vitro transcription: nanoLuc and eGFP coding sequences were amplified by PCR from the pUAS-NanoLuc (Zhang et al. 2017) and pEGFP-N2 plasmids, using primers shown in **Table S2**. Forward and reverse primers were designed to incorporate a T7 RNA polymerase sequence and a Kosak sequence, and a 30-31 nt polyA tail, respectively. PCR reactions were carried out with Phusion Polymerase. The amplification program was: 30 s at 98°C, followed by 30-35 amplification cycles, each comprising 10 s at 98°C, 10 s at 60°C, and 30 s at 72°C. A final extension step of 10 min at 72°C was included. PCR products were analyzed by 1% agarose gel electrophoresis and purified with PureLink quick gel extraction kit directly from the PCR solution (for *nanoLuc*) or from excised gel bands (for *eGFP*). DNA was concentrated by overnight ethanol precipitation and used as template for mRNA synthesis. *nanoLuc* and *eGFP* mRNAs were synthesized and co-transcriptionally capped using HiScribe T7 High Yield RNA Synthesis Kit. Briefly, reactions were performed in a 20-μl reaction volume at 37°C for 2 h using 1 μg of template and a 3’-O-Me-m7G(5’)ppp(5’)G RNA Cap Structure Analog, with 4:1 CAP:GTP ratio, and following manufacturer’s instructions. For *eGFP*, UTP was substituted with N1-Methylpseudouridine-5’-Triphosphate. Afterwards, DNA templates were digested with Turbo DNase I. In vitro-synthesized mRNAs were purified with a Monarch RNA Cleanup Kit and stored at –80°C until use.

### In vitro translation

*NanoLuc* mRNA was translated in vitro with Retic Lysate IVT Kit following manufacturer’s instructions. Briefly, 100 ng of *NanoLuc* mRNA was incubated for 45 min at 30°C with a rabbit reticulocyte extract. Afterwards, nanoLuc bioluminescence was measured using a Bio-Glo-NL Luciferase Assay System and detected in a LUMIstar Optima luminometer.

### nanoLuc mRNA internalization and translation assays

Naked nanoLuc mRNA capture and translation was assessed in murine primary cells (BMDCs) and two human epithelial cell lines (U-2 OS and MCF-7). To quantify *nanoLuc* mRNA levels and nanoLuc protein in BMDCs (**Figure 5B** and **5C**), cells were seeded in 24 (250.000 cells/well) or 48 (120,000 cells/well) well plates, and cultured for 2 h to allow adhesion. Then, growth medium containing FBS was removed, cells were washed, incubated in growth medium without FBS and stimulated for 2 h with 1 μg / mL naked *nanoLuc* mRNA with 80 U / mL RI, thermally-inactivated RI, or without RI. To investigate the impact of varying concentrations of FBS and RI on *nanoLuc* translation by BMDCs (**Figure D**), cells were seeded in 48 well plates (120,000 cells/well) and cultured for 2 h to allow adhesion. Then, growth medium containing FBS was removed, cells were washed, incubated in growth medium containing varying concentration of FBS (0, 0.1, 1, 10%) and RI (0, 80, 120, 240, 320 U / mL) and stimulated with 1 μg / mL naked *nanoLuc* mRNA for 2 h. Subsequently, the medium containing the stimuli was removed and fresh growth medium with 10% FBS was added. After 6 h, *nanoLuc* mRNA was quantified by RT-qPCR. The RT-qPCR protocol was specifically designed to enable detection of full-length, undegraded *nanoLuc* mRNA. This was achieved by designing primers that selectively amplified the transcript 5’ end, which had been previously reverse-transcribed using oligo(dT) primers. After 24 h, nanoLuc protein translation in BMDCs was detected with Bio-Glo-NL Luciferase Assay System using a LUMIstar Optima luminometer. MCF-7 and U-2 OS cells were seeded in 48 well adherent plates. After reaching 80% confluence, growth medium was replaced with growth medium containing 0.1% FBS. Then, cells were incubated with varying concentration of *nanoLuc* mRNA (0, 0.25, 1, 4μg / mL), with either 80 U / mL RI or thermally inactivated RI for 2 h. To assess the involvement of dynamin-dependent endocytosis in naked *nanoLuc* mRNA capture, U-2 OS cells were incubated for 2 h with 80 μM Dynasore or DMSO, before addition of 1 μg / mL *nanoLuc* mRNA, with or without 80 U / mL RI. After 2 h, media was replaced with growth medium containing 10% FBS. Cells were cultured for additional 24 h, and nanoLuc protein levels were measured as previously described.

### naked eGFP internalization and translation assays

U-2 OS cells were seeded onto 35-mm glass bottom dishes (100,000 cells) and incubated overnight with growth medium. The next day, cells were washed 3 times with MEGM (Lonza), and MEGM either without or with 0.1% FBS was added. Cells were incubated for 2 h with 5 μg / mL naked eGFP mRNA in the presence or absence of 80 U / mL RI, or with a DOTAP-eGFP mRNA complex. After the incubation, cells were washed 3 times and cultured for an additional 24 h in complete growth medium, containing 10% FBS. To visualize and analyze eGFP translation, cells were imaged using a Zeiss LSM 880 confocal microscope. Nuclei were stained with NucRed (Thermo). Imaging was performed utilizing a 63x oil immersion objective with a numerical aperture of 1.4, employing a Plan Apochromat configuration. The imaging acquisition parameters were set with a pixel dwell time of 4.10 microseconds and size of 0.55 microns, utilizing bidirectional scanning. The total scanning duration for each acquisition was 1 minute and 20 seconds, with an image size of 655.8 x 655.8 microns. Images were subsequently processed and analyzed using ImageJ. The segmentation was conducted on both the green (eGFP) and red (nuclei) fluorescence channels. Quantitative analysis of the cellular responses was executed through Integrated Density calculations. Prior to calculations, a noise subtraction step was implemented to ensure data integrity. The resultant processed images were subjected to total image calculations, facilitating the extraction of Integrated Density values indicative of cellular responses.

### In vivo experiments

For intraperitoneal administration of RNAs, female mice were *i.p* inoculated with 250 µl of varying doses (0.025 – 62.5 µg) of naked total *E. coli* RNA with or without 40, 200 or 400 units of RI in DPBS. DPBS and 62.5 µg LPS were used as positive and negative controls, respectively. After 24 h, animals were euthanized and 4-5 ml of DPBS supplemented with 3 mM EDTA was injected into the peritoneal cavity to extract peritoneal cells. Peritoneal cells were pelleted at 375 g for 5 min at 4°C, resuspended in FACS buffer (0.5% w/v BSA, 2 mM EDTA in DPBS), counted, and stained for flow cytometry analysis. For systemic administration of RNAs, female mice were retro-orbitally inoculated with 100 µl of either high molecular weight naked poly(I:C) (200 ng or 1000 ng) or naked total *E. coli* RNA 10 µg in the presence or absence of 480 units of RI in DPBS. DPBS with or without RI was used as control. After 6 h, mice were euthanized, and spleens were collected. To obtain spleen cells, spleens were incubated in 1ml of collagenase solution (2 mg / mL Collagenase D, 2% v/v FBS in RPMI), cut on ice with a bistoury and incubated for 20 min at 37°C. The reaction was stopped with 100 µl of 100 mM EDTA. Spleen cells were passed through a 100 µm filter and resuspended in 40 ml of PES buffer (0.7 mM EDTA, 2% v/v FBS in DPBS). Cells were pelleted at 1500 rpm for 15 min at 4°C, resuspended in 5 ml RBCL buffer (154 mM NH_4_Cl, 10 mM KHCO_3_, 0.1 mM EDTA; pH = 7.2-7.4), and incubated for 10 min at room temperature to lyse red blood cells. Then, spleen cells were washed two times with PES solution, resuspended in FACS buffer, counted, and stained for flow cytometry analysis.

### Flow cytometry

Cells suspended in FACs buffer were seeded in V-shaped 96 well plates (200,000-500,000 cells/well). When live/dead viability dye was used, cells were stained with 200 µl of 1/200 live/dead violet in FACS buffer and incubated on ice for 30 min in the dark. If viability was assessed with DAPI, the previous step was not performed. Then, cells were pelleted at 2200 rpm for 2 min and blocked with 25 µl of 10% normal rat serum in FACS buffer for 30 min on ice. For cell surface antibody staining, 25 µl of 2x antibody mix (see **Table S3**) in FACS buffer was added, and cells were stained for 30 min on ice in the dark. Stained cells were washed 2 times with FACS buffer, resuspended in 200 µl FACS buffer, and analyzed by flow cytometry. To assess viability, 0.5 μg / mL final concentration of DAPI was added immediately before sample analysis. Myeloid and lymphoid spleen cells were analyzed in a Cytek Aurora spectral flow cytometer. Peritoneal immune cells were analyzed in an Attune NxT flow cytometer or in a Cytek Aurora spectral flow cytometer. BMDCs cells were analyzed in a BD Accuri C6 flow cytometer or in a Cytek Aurora spectral flow cytometer.

### RNA Polyacrylamide electrophoresis

RNAs were analyzed by polyacrylamide gel electrophoresis under denaturing conditions using 6-10% acrylamide TBE-urea gels. Briefly, RNA samples were mixed with 2x RNA loading dye, heated at 65°C for 5 min, cooled on ice and analyzed on an XCell *SureLoc* Mini-Cell Electrophoresis system (Invitrogen) using 0.5x TBE as running buffer. Gels were run at constant voltage (160V) for 60-80 min. Gels were stained for 15 min with 1/10000 SYBR Gold in 0.5x TBE, and imaged with a transilluminator (Loccus biotecnologia) with a coupled imaging platform (Carestream). When needed, images were cut and contrasted using PhotoScapeX.

## QUANTIFICATION AND STATISTICAL ANALYSIS

Statistical analysis was performed using R or GraphPad Prism 9. For RNA-seq experiments, 3 biological replicates for all 4 experimental conditions were performed. Significant differences in gene expression across samples was computed with DESeq2 and multiple hypothesis testing was controlled for using the Benjamini-Hochberg method to estimate false discovery rates (FDR). Differentially expressed genes with adjusted p-values lower than 0.001 were reported. For in vitro experiments each dot in bar plots represents a biological replicate. For in vivo experiments, each dot in bar plots represents a different mouse. For microscopy experiments, each dot in violin plots represents an individual cell. When statistical analysis was performed, normal distribution of data were assessed using Shapiro-Wilk and Kolmogorov-Smirnov tests. If data was normally distributed and had variance homogeneity, one-way ANOVA with multiple comparisons and Fischer’s LSD test was performed. If data was normally distributed but had no variance homogeneity, Welch ANOVA tests was employed with multiple comparisons and unpaired Welch’s correction test. If data had no normal distribution, Kruskal-Wallis test with multiple comparisons and Dunn’s post-test was performed. Statistically significant differences are indicated with asterisks (*: p-value < 0.05, **: p-value < 0.005, ***: p-value < 0.0005, ****: p-value < 0.0001).

## SUPPLEMENTARY FIGURES

See Supplementary Figures file for Fig S1 – S10

## SUPPLEMENTARY TABLES

**Table S1.**
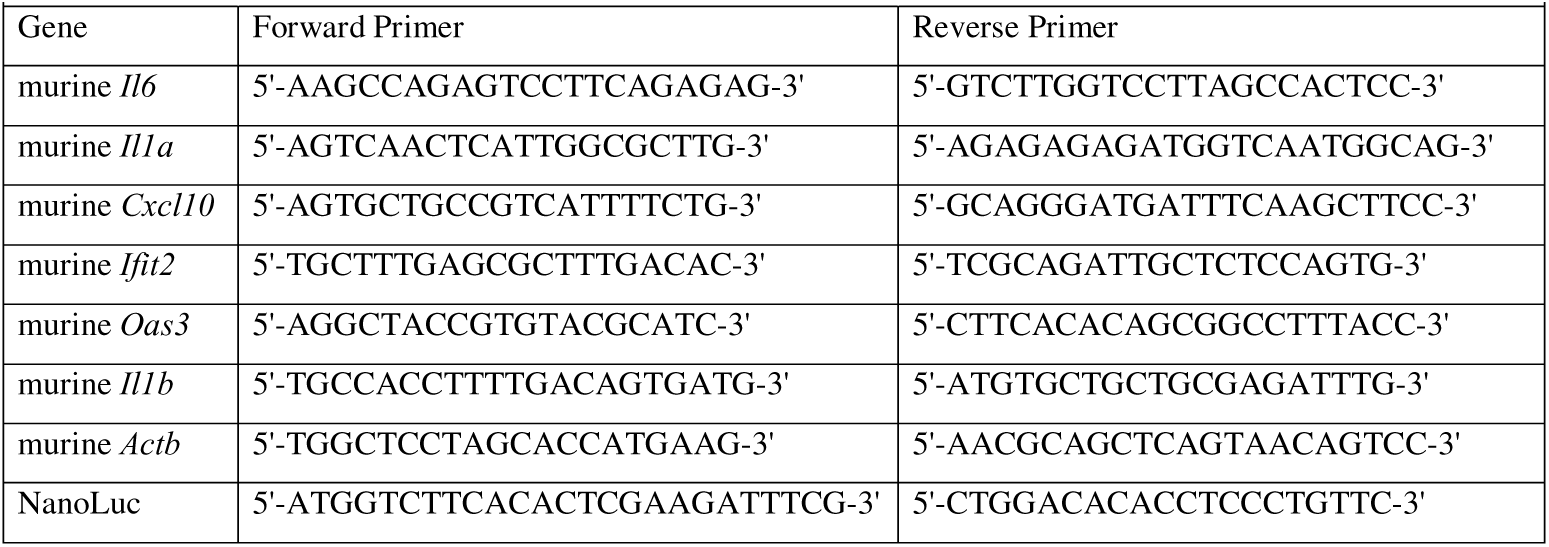
Primers for RT-qPCR.

**Table S2.**
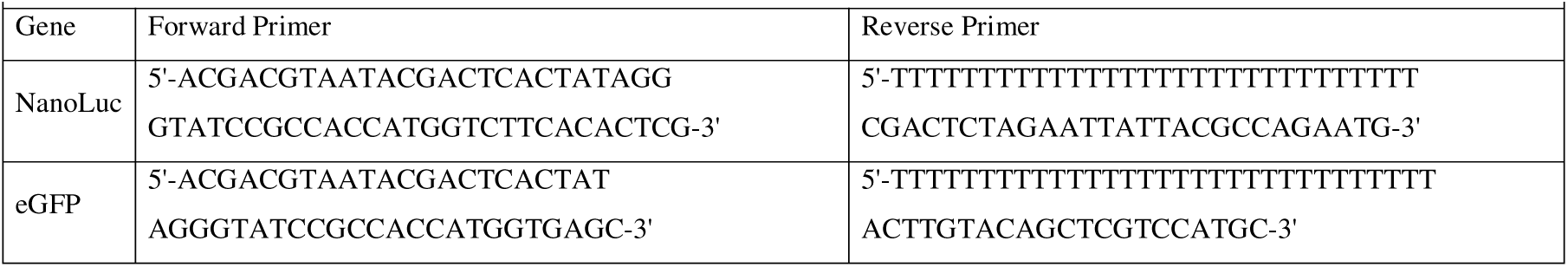
Primers for IVT template amplification.

**Table S3.** Antibodies mixes employed.

| . | Antibodies | Concentration |
| --- | --- | --- |
| BMDCs | CD40-FITC | 1/500 |
|  | CD86-PE | 1/200 |
|  | MHCII-APC | 1/200 |
| Spleen myeloid cells | CD19-APC-Cy7 | 1/200 |
|  | TCR-B-APC-Cy7 | 1/200 |
|  | Ly6G-BV711 | 1/200 |
|  | CD11c PE-Cy7 | 1/200 |
|  | MHC-II-APC | 1/200 |
|  | B220-PerCP-Cy5.5 | 1/200 |
|  | F4/80-PE | 1/200 |
|  | CD11b-AF700 | 1/200 |
|  | CD86-FITC | 1/500 |
| Spleen lymphoid cells | CD19-PerCP-Cy5.5 | 1/200 |
|  | TCR-B-APC-Cy7 | 1/200 |
|  | CD69-PECy7 | 1/200 |
|  | CD86-FITC | 1/500 |
| Peritoneal macrophages and monocytes | CD19-APC-Cy7 | 1/200 |
|  | TCR-B-APC-Cy7 | 1/200 |
|  | CD11b-AF700 | 1/200 |
|  | F4/80-PE | 1/200 |
|  | Ly6c-FITC | 1/500 |

