## Supplementary Figures file for "Ribonuclease activity undermines immune sensing of naked extracellular RNA"

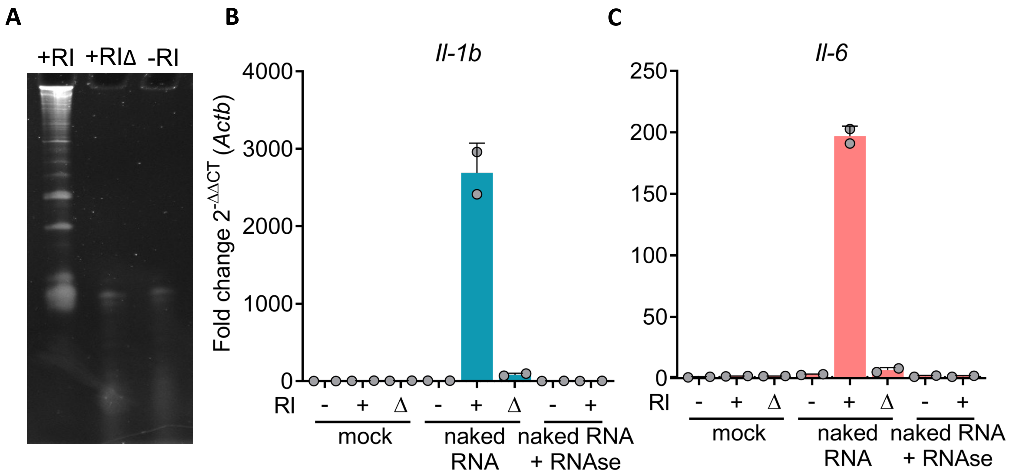


**Supplementary Figure 1. Inflammatory naked extracellular RNA recognition is completely dependent on RI enzymatic activity.**

**A)** PAGE under denaturing conditions of U-2 OS total RNA incubated in 10% fetal bovine serum together with 400 U / mL RI (+RI) or heat inactivated RI (+RIΔ) or without RI (-RI) for 45 minutes. **B,C)** *Il-1b* **(B)** and *il-6* **(C)** expression by RT-qPCR in BMDCs stimulated for 6 hs with 1 µg / mL naked total RNA from *E. coli* with RI 80 U / mL (+RI) or heat inactivated RI (+RIΔ) or without RI (-). RNAse1 treated *E. coli* RNA with (+RI) or without RI (-) were used as control and DPBS as negative control.

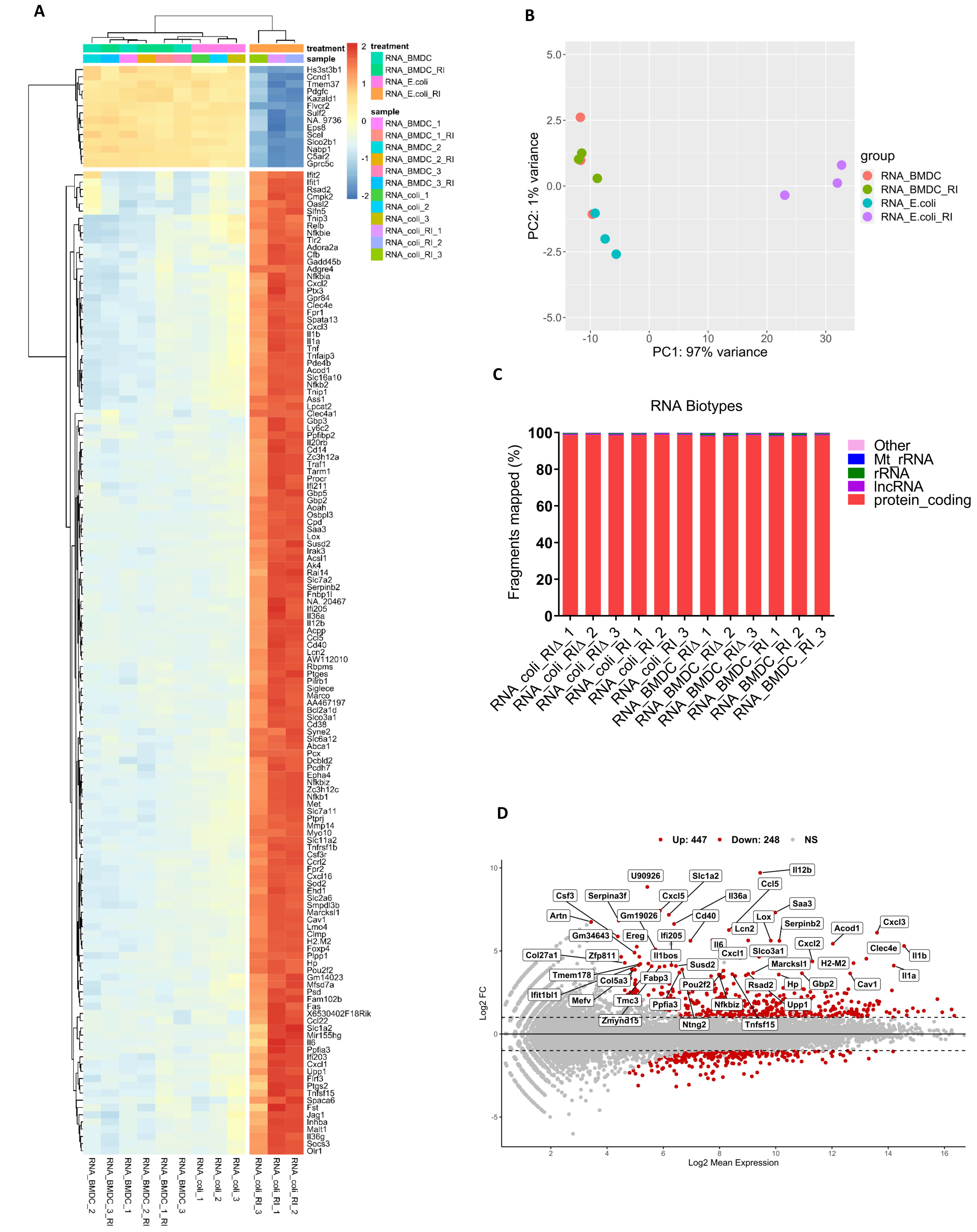


**Supplementary Figure 2**

**A)** Heatmap showing top 150 most differentially regulated genes in BMDCs stimulated for 6hs with 100ng/ml naked total RNA from *E. coli* or 100ng/ml total RNA from BMDCs in the presence of RI 80 U/ml (+RI) or thermally inactivated RI (+RIΔ). **B)** PCA plots corresponding to all 12 experimental conditions. **C)** RNA biotypes abundance across samples. **D)** MA-plot between BMDCs stimulated with 100ng/ml naked total RNA with RIΔ or RI.

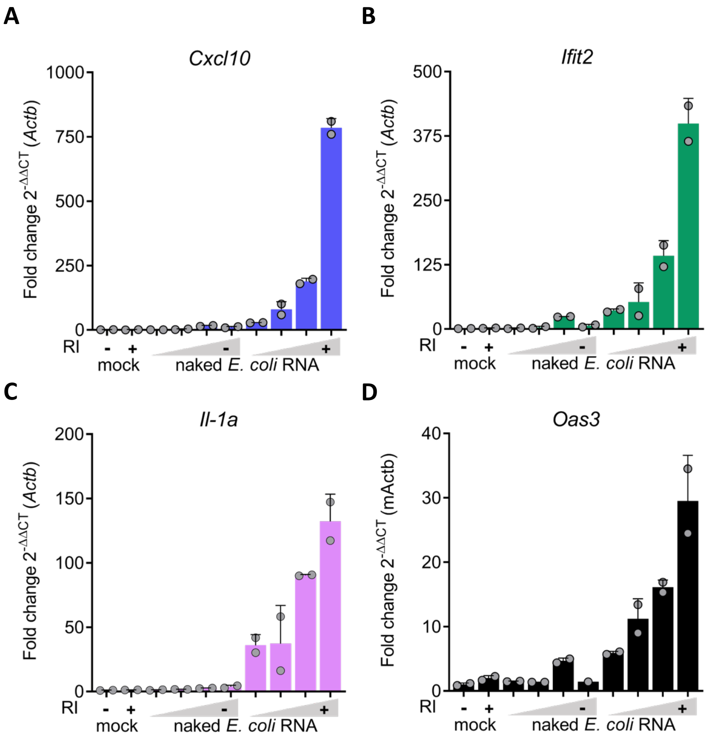


**Supplementary Figure 3**

*Cxcl10* **(A)** *ifit2* **(B)** *Il-1a* **(C)** *and Oas3* **(D)** expression by RT-qPCR in BMDCs stimulated for 6hs with varying doses (1; 5; 10; 25 µg / mL) of naked total RNA with or without RI 80 U / mL. DPBS was used as negative control.

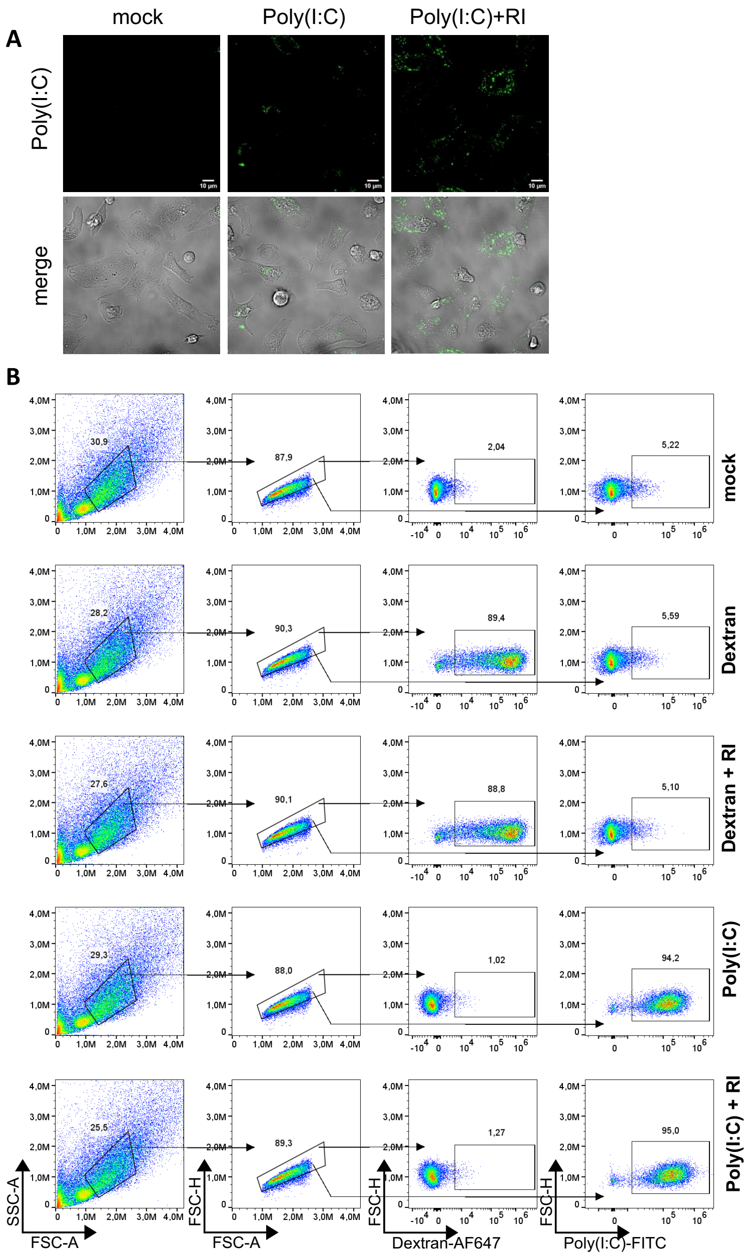


**Supplementary Figure 4. Internalization of naked fluorescent RNA by BMDCs**

**A)** Confocal microscopy of BMDCs cultured in 10% FBS left untreated or stimulated with naked Poly(I:C)-Fluorescein 0.5 µg / mL with or without RI for 1 hr. **B)** Flow cytometry of BMDC cultured in 10% FBS and stimulated for 1 hr with naked Poly(I:C)-Fluorescein 0.5 µg / mL or Dextran-AF647 with or without RI 120 U / mL.

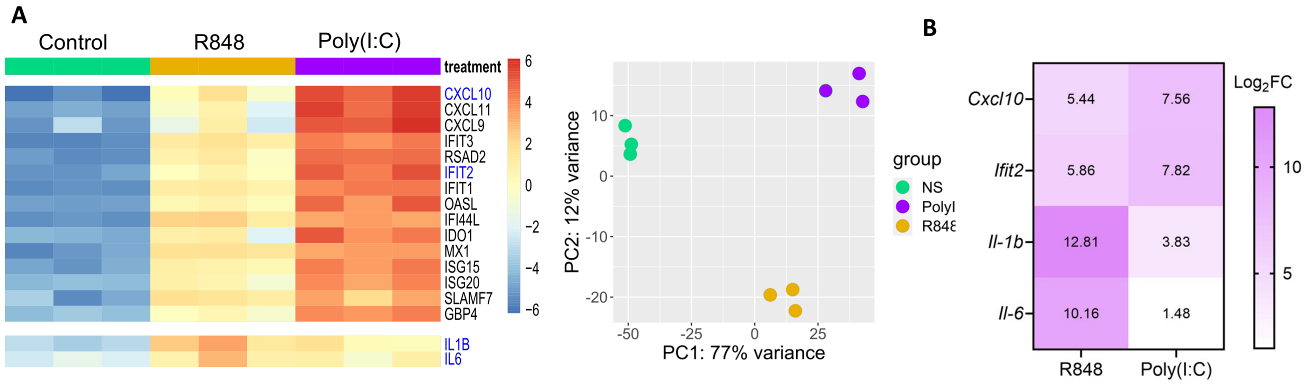


**Supplementary Figure 5**

**A)** Data mining and differential gene expression analysis of a transcriptomic dataset (PMID:31968263, GEO:GSE125817) from monocyte derived dendritic cells stimulated with Poly(I:C) or R848 for 6 h. A heatmap with the top 15 most upregulated genes (including *Il-1b* and *Il-6*) and the corresponding PCA plot are shown. **B)** Heatmap showing *Cxcl10, Ifit2, Il-1b* and *il-6* expression by RT-qPCR in BMDCs stimulated for 6hs with 100 ng / mL R848 or 10 µg / mL poly(I:C). DPBS was used as negative control.

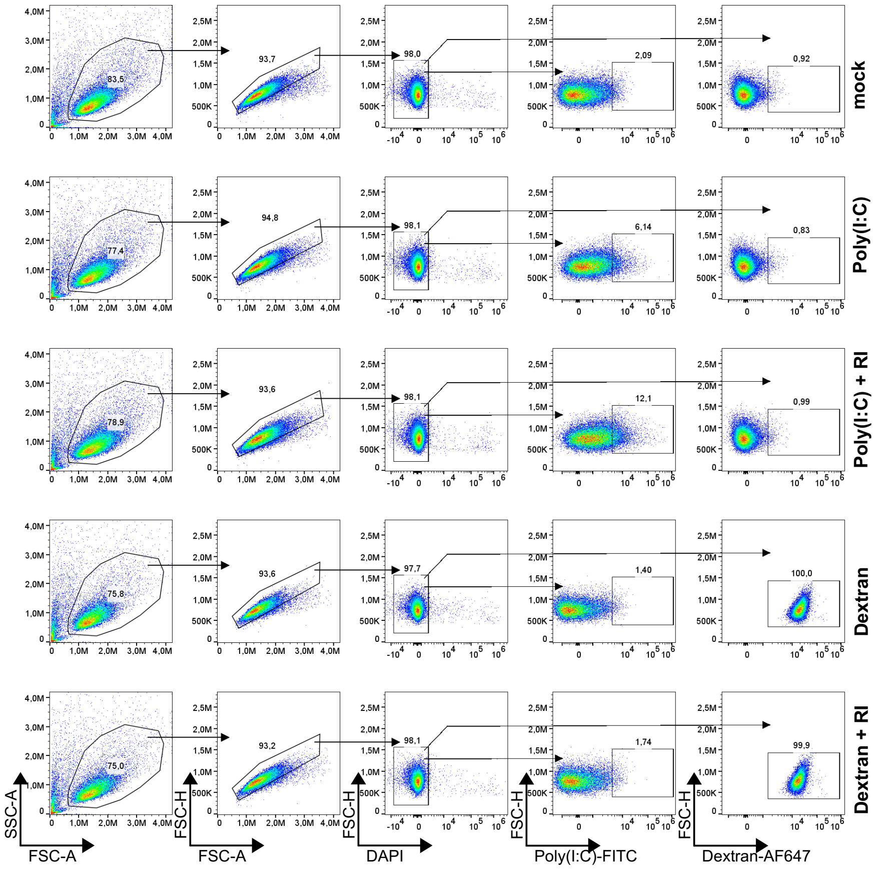


**Supplementary Figure 6. Internalization of naked fluorescent RNA by THP1 cells**

Flow cytometry of THP1 cells cultured in 10% FBS and stimulated for 1 h with naked Poly(I:C)-Fluorescein 0.5 µg / mL or Dextran-AF647 with or without RI 120 U / mL.

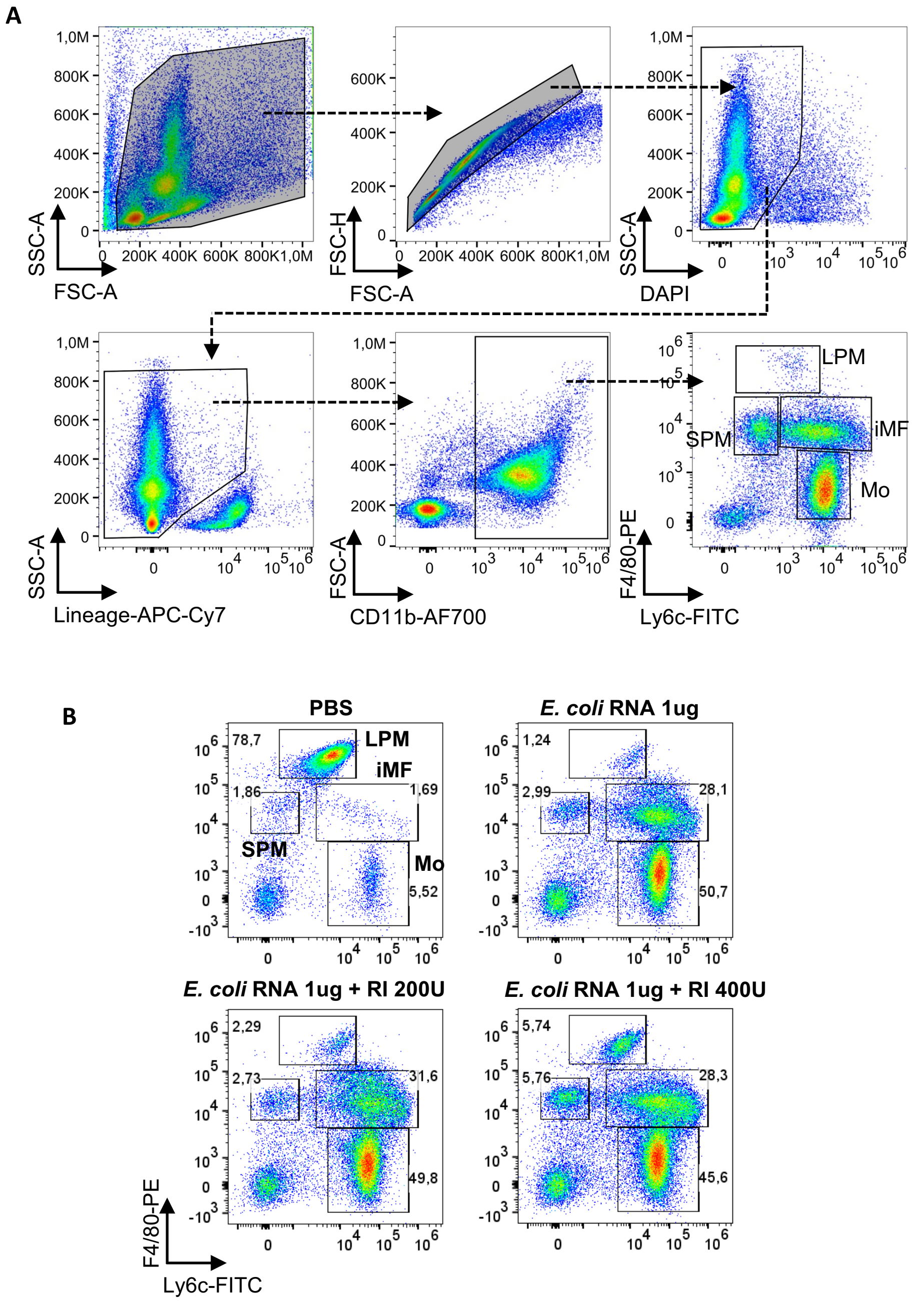


**Supplementary Figure 7. Gating strategy to study of mice peritoneal cells**

Flow cytometry gating strategy to analyze resident large (LPM) and small (SPM) peritoneal macrophages, infiltrating monocytes (Mo) and inflammatory macrophages (iMF). Gating strategy shows cells 24 h after *i.p* administration of 12,5 µg of total naked *E. coli* RNA **(A)** Peritoneal cell populations gated under CD11b+Lineage- 24hs after administration of PBS or naked *E. coli* RNA 1 µg with the indicated RI dose **(B).** Percentages correspond to the parental gates.

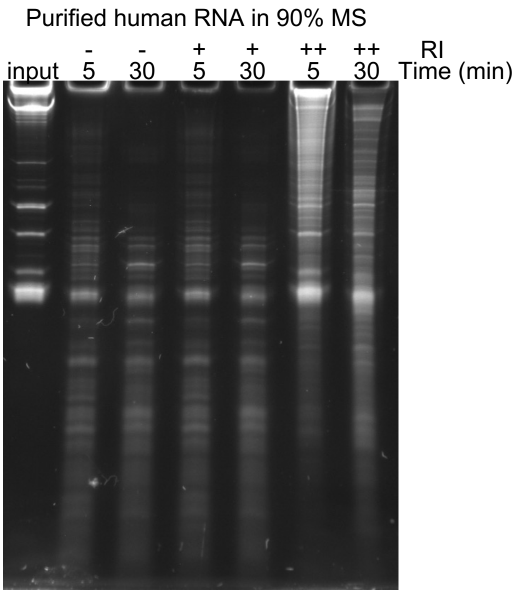


**Supplementary Figure 8.**

PAGE under denaturing conditions of human total RNA incubated in 90% mouse serum for either 5 or 30 minutes in the presence of 160(++), 40(+) U / mL of RI or no RI(-).

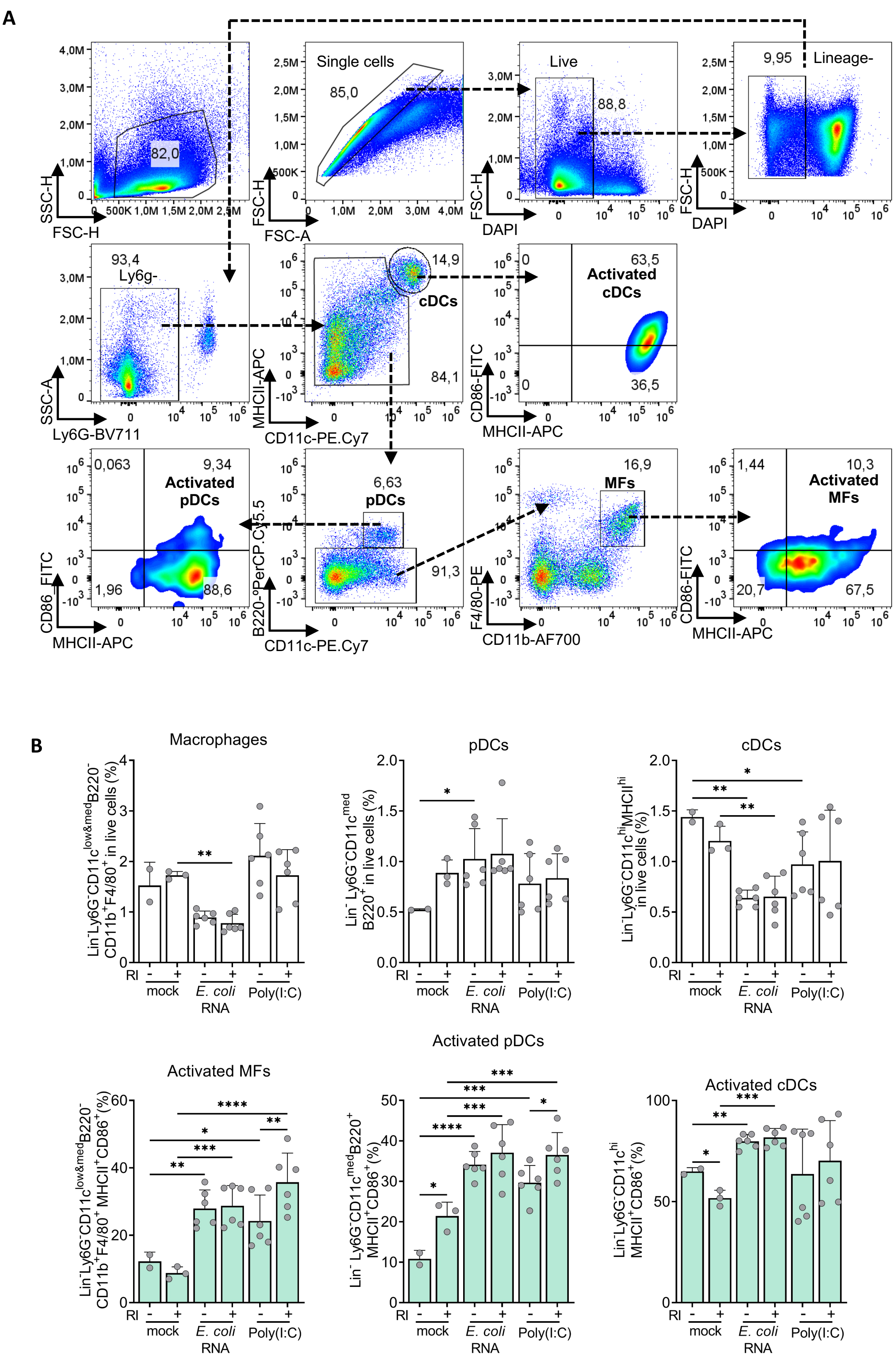


**Supplementary Figure 9. Effect of naked exRNA in vivo on splenic myeloid cells and gating strategy**

**A)** Flow cytometry gating strategy to study splenic cDC, pDC and macrophages after intravenous administration of naked RNA stimuli. Percentages correspond to the parental gates. **B)** Flow cytometry analysis of spleen cells 6 h after intravenous administration of either naked Poly(I:C) 200ng, naked total *E. coli* RNA 10 µg with or without 480 units of RI. PBS was used as mock. The percentage of total Macrophages (Lin$^-$Ly6G$^-$CD11c$^{low\&med}$B220$^-$CD11b$^+$F4/80$^+$), pDCs (Lin$^-$ Ly6G$^-$CD11c$^{med}$B220$^+$), cDCs (Lin$^-$Ly6G$^-$CD11c$^{hi}$MHCII$^{hi}$), gated in live cells is shown in top panels. The percentage of activated macrophages, pDCs, cDCs, (defined as MHCII$^+$CD86$^+$) gated under their corresponding cell type is shown in bottom panels with bar plots shaded in green.

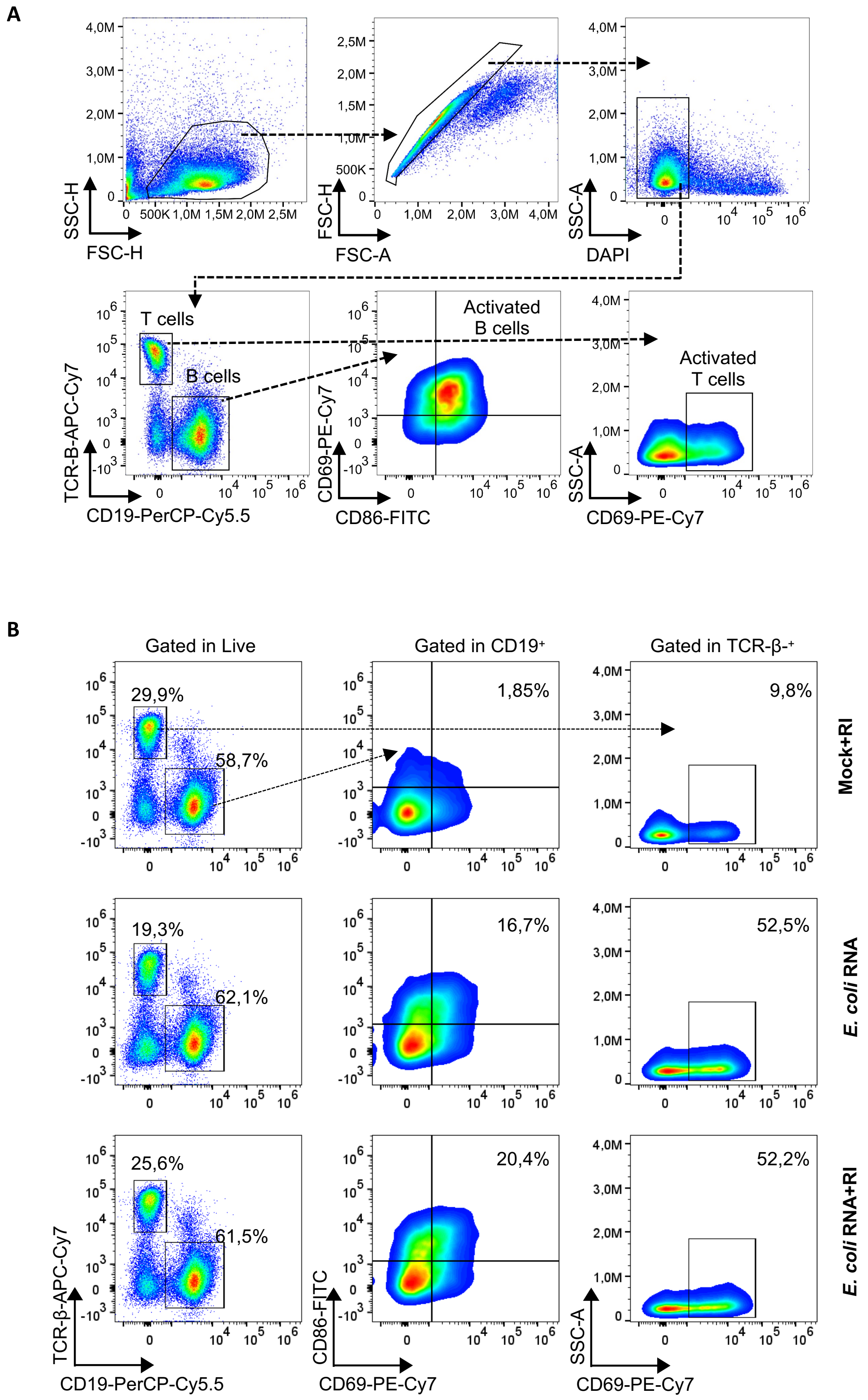


**Supplementary Figure 10. Effect of naked exRNA in vivo on splenic lymphoid cells and gating strategy**

**A)** Flow cytometry gating strategy to study splenic B and T cells populations after intravenous administration of RNA stimuli. Percentages correspond to the parental gates. **B)** Dot plots showing flow cytometry analysis of spleen cells 6 h after intravenous administration of naked total *E. coli* RNA 10 µg with or without 480 units of RI and PBS with RI. Percentages correspond to the parental gates.
